# Inflammation in the tumor-adjacent lung as a predictor of clinical outcome in lung adenocarcinoma

**DOI:** 10.1101/2022.11.10.516003

**Authors:** Igor Dolgalev, Hua Zhou, Nina Shenker-Tauris, Hortense Le, Theodore Sakellaropoulos, Nicolas Coudray, Kelsey Zhu, Varshini Vasudevaraja, Anna Yeaton, Chandra V. Goparaju, Yonghua Li, Imran Sulaiman, Jun-Chieh J. Tsay, Peter Meyn, Hussein Mohamed, Iris Sydney, Sitharam Ramaswami, Navneet Narula, Ruth Kulicke, Fred P. Davis, Nicolas Stransky, Gromoslaw A. Smolen, Wei-Yi Cheng, James Cai, Salman Punekar, Vamsidhar Velcheti, J.T. Poirier, Ben Neel, Kwok-Kin Wong, Luis Chiriboga, Adriana Heguy, Thales Papagiannakopoulos, Matija Snuderl, Leopoldo N. Segal, Andre L. Moreira, Harvey I. Pass, Aristotelis Tsirigos

## Abstract

Early-stage lung adenocarcinoma is typically treated by surgical resection of the tumor. While in the majority of cases surgery can lead to cure, approximately 30% of patients progress. Despite intense efforts to map the genetic landscape of early-stage lung tumors, there has been limited success in discovering accurate biomarkers that can predict clinical outcomes. Meanwhile, the role of the tumor-adjacent tissue in cancer progression has been largely ignored. To test whether tumor-adjacent tissue can be informative of progression-free survival and to probe the underlying molecular pathways involved, we designed a multi-omic study in both tumor and matched tumor-adjacent histologically normal lung tissue from the same patient. Our study includes 143 treatment naive stage I cases with long-term patient follow-up and is, to our knowledge, the largest such study with the longest follow-up. We performed a comprehensive histologic characterization of all tumors, mapped the mutational landscape and probed the transcriptome of both tumor and adjacent normal tissue. We evaluated the predictive power of each data modality and showed that the transcriptome of tumor-adjacent histologically normal lung tissue is the only reliable predictor of clinical outcome. Unbiased discovery of co-expressed gene modules revealed that inflammatory pathways are upregulated in the tumor-adjacent tissue of patients at high risk for disease progression. Furthermore, single-cell transcriptome analysis in the tumor-adjacent lung demonstrated that progression-associated inflammatory signatures were broadly expressed by both immune and non-immune cells including mesothelial cells, alveolar type 2 cells and fibroblasts, CD1 dendritic cells and MAST cells. Collectively, our studies suggest that molecular profiling of tumor-adjacent tissue can identify patients that are at high risk for disease progression.

## Introduction

Despite advances in diagnosis and treatment, lung adenocarcinoma (LUAD), the most prevalent non-small cell cancer, remains the deadliest cancer in the United States. The risk of disease progression for early non-small cell lung cancer patients is currently about 30% after surgery^1^. With the emergence of improved treatments, recent studies have focused on creating predictive models for progression-free survival (PFS) and overall survival (OS) in lung cancer based on histology, mutations, gene expression, proteomics, and microbiome. Several studies have analyzed correlations between prognosis in resected early stage LUAD patients and histopathological patterns such as histological grading, predominant and high-grade patterns^2, 3^, and quantitative morphological features from histopathological images extracted with machine learning algorithms^4, 5^. However, studies integrating histology lack validation in clinical settings. Gene mutations in SMARCA4 and TP53^6^, in ATR, ERBB3, KDR, and MUC6^7^, and fusions in GOPC-ROS1 and NTRK1-SH2DA^7^ have also been identified as potential biomarkers for early stage LUAD recurrence after surgical resection, in contrast with EGFR which does not impact survival in early stages^8^. Nevertheless, these mutation prognostic tools need to be tested on independent external datasets. Gene expression is currently a growing field for the discovery of clinically relevant biomarkers for lung cancer recurrence prediction. Diverse machine learning algorithms integrated gene expression signatures and gene-expression based molecular subtypes, and selected key genes to elaborate prognostic models for lung cancer^9, 10^. However, these studies lack clinical reproducibility. Proteomics biomarkers have also been the center of many current studies. Models integrating proteins with distinct proteomic changes^11^, or incorporating a proteomics score^12^ were correlated to survival in NSCLC although they need to be validated on independent large-scale datasets as well. The emphasis thus far in studies attempting to stratify early-stage lung cancer has concentrated on signatures from the tumor itself. In this study, we hypothesized that tumor-adjacent normal lung tissue may hold significant prognostic information in early-stage lung cancer. Although a few studies suggested that airway transcriptomic profiles could add value to bronchoscopy for the diagnosis of the indeterminate pulmonary nodule without a direct biopsy of the tumor^13–20^, a review of the literature reveals that there have been no studies for the prognostication of lung cancer which investigate the transcriptome of lower airway samples, specifically using matched lung tissue from lung cancer bearing individuals. To investigate whether the transcriptome of the tumor-adjacent normal tissue can predict disease progression, we designed a matched tumor-normal study of early-stage lung adenocarcinoma patients with extensive follow-up. We used DNA-sequencing and RNA-sequencing to map the mutational and transcriptomic landscape respectively in this cohort in both the tumor and the tumor-adjacent normal. Our data shows that the transcriptome obtained from normal lung tissue, rather than that of the tumor, is the best predictor of progression. Furthermore, using unsupervised clustering for the *de novo* unbiased discovery of co-expressed gene modules, we identified a gene module characterized by TNF-α, NFκB and IL-17 signaling which is uniquely activated in the tumor-adjacent normal tissue of patients that eventually progress. We show that a simple inflammatory score derived without supervised training and/or a complicated set of parameters can effectively stratify patients by risk of disease progression. Using public datasets from TCGA, we show that the same inflammatory score can stratify patients in other cancer types. Finally, using single-nucleus RNA-sequencing on a subset of samples from our cohort, we discovered the cell types that are the main source of the prognostic inflammatory signature.

## Results

### A matched tumor-normal lung study: design and cohort characteristics

In this study, we used a treatment-naive stage I lung adenocarcinoma cohort of patients with matched tumor and tumor-adjacent histologically normal lung samples (within the same lobe, segment, or wedge resection) obtained from our biorepository of prospectively collected specimens (see Methods). Subjects who received antibiotics (except peri-operative), steroids, radiation, immunotherapy, or chemotherapy within the month prior to surgery were excluded from the study. A total of 143 patients matched our inclusion and exclusion criteria (**Figure 1a**). To our knowledge, this is the largest study of matched tumor-normal early-stage cancer, as TCGA is limited to only 54 stage I patients with matched tumor-normal samples (**Figure 1b**). Detailed information about our cohort and a comparison with the TCGA stage I cohort can be found in **Supplementary File 1**. Overall, there were no major differences between the two cohorts in terms of patient characteristics, although our cohort consisted of slightly older patients (**Supplementary Figure 1a**) with a lower median of pack years (**Supplementary Figure 1b**).

**Figure 1.**
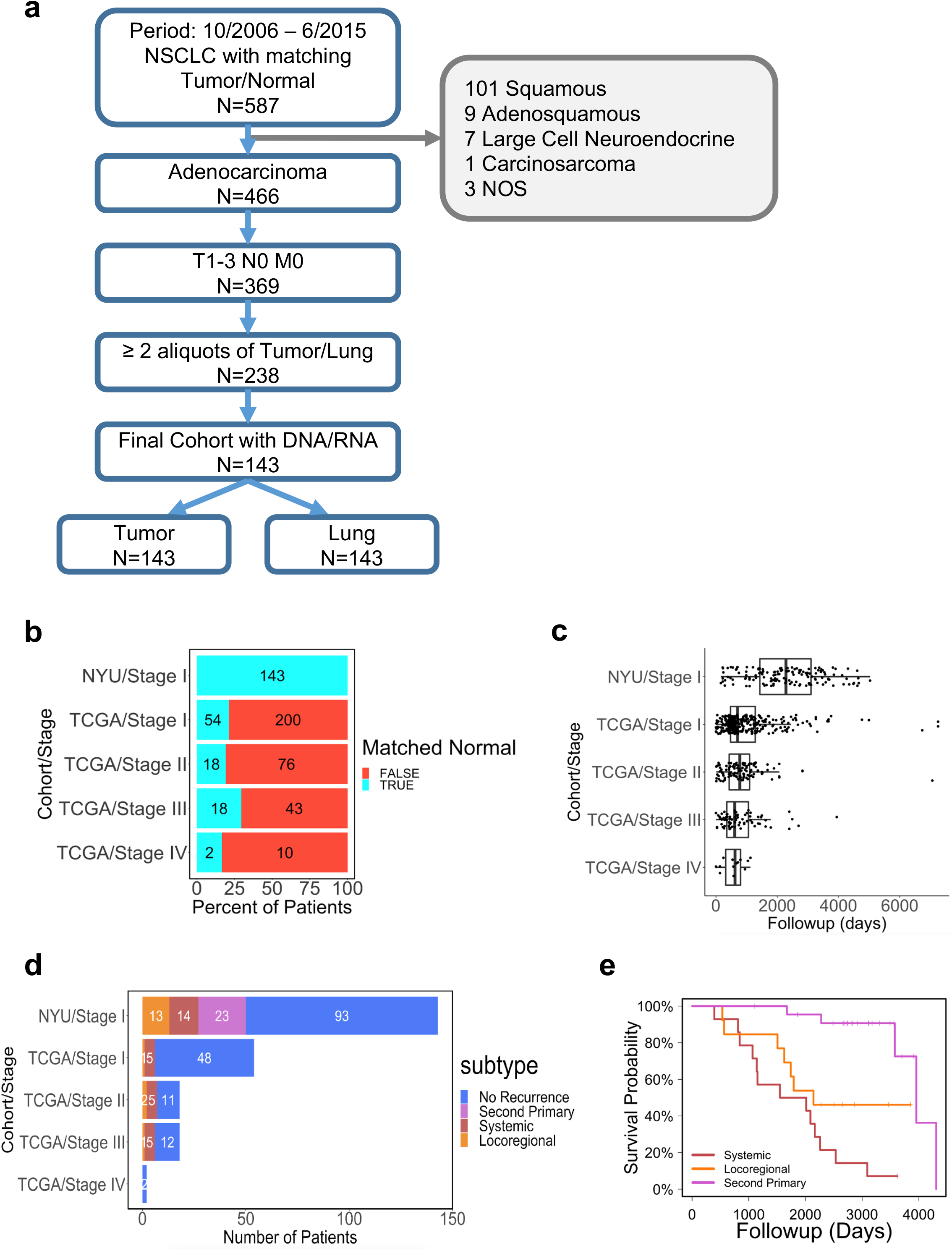
Study design and cohort characteristics. (a) CONSORT diagram, (b) Availability of matched tumor-adjacent normal lung patient samples in the NYU and TCGA cohorts, (c) Patient follow-up distribution in NYU Stage I cohort and in stage-specific TCGA cohorts, (d) Number of patients with available matched normal lung samples by progression type across the NYU and TCGA cohorts, (e) Overall survival (OS) of patients with recurrence (systemic, locoregional) or second primary tumors.

Importantly, our cohort has an extensive follow-up, while the follow-up time in TCGA is rather limited (median follow-up of 2,284 days versus 701 days) (**Figure 1c**). Substantially longer follow-up allows us to observe a significant number of disease progression events and enable the discovery of molecular signatures of progression-free survival. To date, we have recorded 50 patients (35%) in our cohort with disease progression. Specifically, we have identified 23 patients who developed a second primary tumor in the lung, 13 patients have been diagnosed with locoregional recurrence in lymph nodes or tumor bed, and 14 with systemic metastasis in the brain, bone, pleura, liver, or adrenal gland; by comparison, only 6 patients have been documented with progressed disease in the TCGA stage I cohort (**Figure 1d**). Distributions of age, smoking, sex, histologic and International Association for the Study of Lung Cancer (IASLC) grade in the progression and no progression groups are shown in **Supplementary Figures 1c-g**. Univariate Cox regression analysis recapitulated previous results (**Supplementary Figure 2**). The overall survival for patients with systemic or locoregional recurrence is worse than in patients with a second primary tumor (**Figure 1e**).

### Mutational and transcriptomic profiling of matched tumor-normal lung specimens

We first performed DNA sequencing of the patient samples using the NYU GenomePACT panel which covers the exons of 580 protein-coding genes plus the TERT promoter (see Methods for details). For each patient, we used samples from the tumor, adjacent normal lung and normal blood (see **Supplementary File 2** for quality assessment). We then performed RNA-seq on all 286 samples (143 tumors and 143 tumor-adjacent normal lungs). The RNA-seq analysis generated adequate sequenced reads and high percentages of uniquely aligned reads for the majority of samples (**Supplementary File 3**): 15 tumor and 10 normal lung samples were excluded from downstream analysis due to low library quality. Eventually, 123 matched tumor-normal samples (86% of the initial set of 143 matched samples) were deemed high-quality RNA-seq samples and used for the downstream analyses. As expected, Principal Component Analysis (PCA) of the RNA-seq data shows a separation of tumor and normal samples (**Supplementary Figure 3a**). **Supplementary Figure 3b** summarizes the workflow of sequencing and quality control. In summary, the vast majority of the patients in our cohort successfully underwent mutational and transcriptional profiling using a total of 5 samples per patient. To ensure that each set of 5 samples indeed belongs to the same patient and exclude possibility of sample swapping and/or mislabeling during sample collection, library preparation or sequencing, we performed a relatedness analysis based on common variants (see Methods for details). The full results of the genotyping analysis are included in **Supplementary Figure 4** and demonstrate that the different samples were all properly labeled.

### Mutations are poor predictors of clinical outcome in early-stage lung adenocarcinoma

Analysis of the DNA sequencing data obtained from the patients’ tumors revealed a mutational landscape with the typical distribution of frequently mutated genes in early-stage lung adenocarcinomas: 34% EGFR, 25% KRAS and 7% STK11 (**Figure 2a**). We then looked at genes that may be mutated at different rates in patients that progress compared to those that do not. We defined two groups, the progression group comprising all disease progression events regardless of the progression type and the no progression group comprising all patients that did not progress with at least 5-year follow-up. Of the most frequently mutated genes, EGFR and TP53 are found in the same percentage of patients in both groups, while KRAS and STK11 are found more frequently in the progression group (**Supplementary Figure 5a**). As expected, stratifying patients by EGFR mutational status does not yield a statistical difference in PFS (**Supplementary Figure 5b**), while even a stratification by KRAS or STK11 mutational status is not significant (p-value>0.01, **Figure 2b,c**). We next tested tumor mutational burden (TMB) as a predictor of disease progression: predicting 5-year recurrence by using TMB for patient ranking yielded an AUC of only 0.62 (**Figure 2d**). We also called mutations in the tumor-adjacent normal samples (using normal blood as control) and found that the driver mutations identified in the tumor samples were not detected in the matched tumor-adjacent normal samples, providing evidence that there is no detectable mutations in the normal samples, which would indicate a potential contamination with tumor cells.

**Figure 2:**
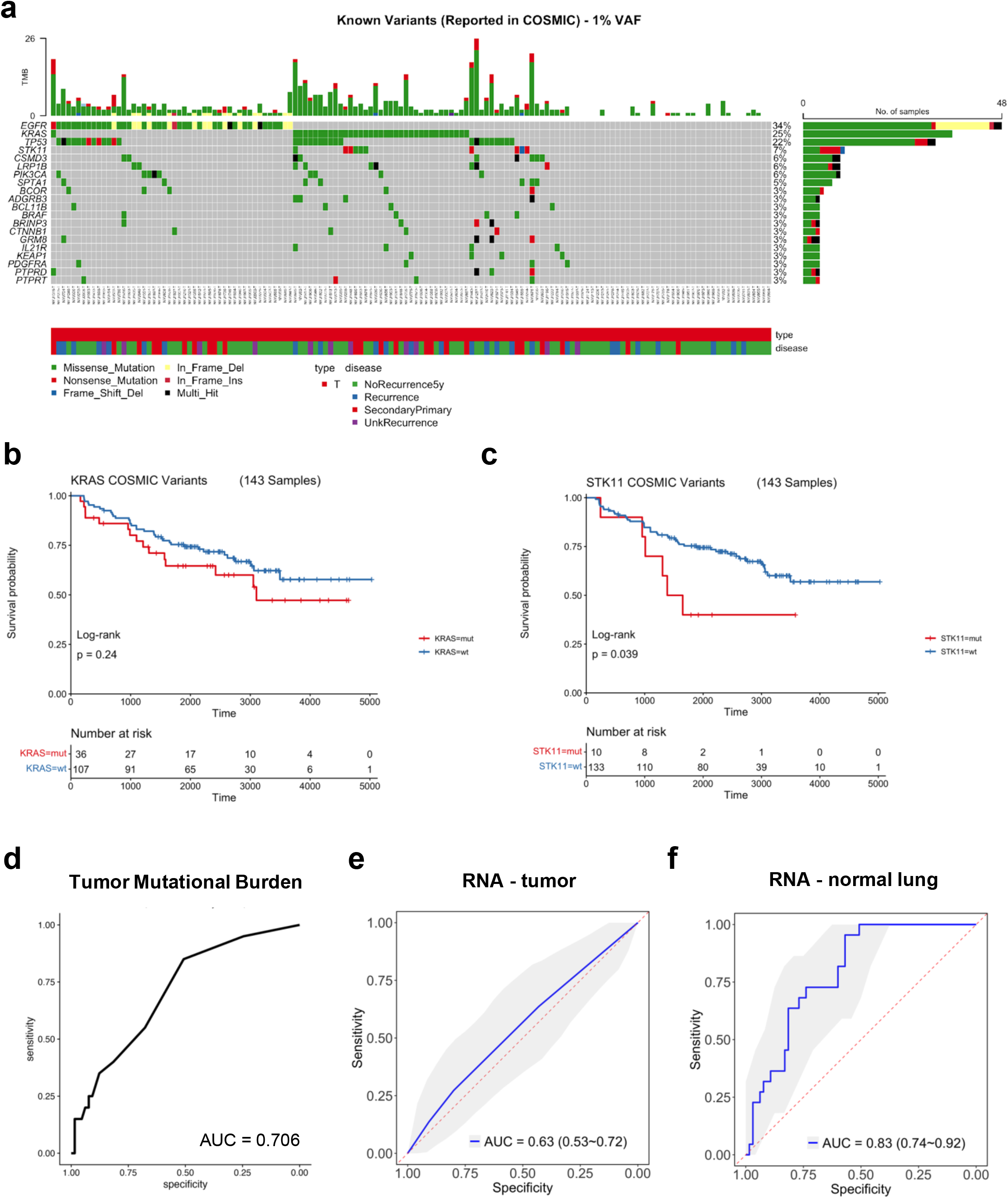
Multi-omic profiling of matched tumor-normal stage I lung adenocarcinomas. (a) Oncoprint of frequently mutated genes in the tumor samples (type T stands for tumor) (b) Kaplan-Meier progression-free survival (K-M PFS) plots comparing patients with and without KRAS mutation (c) K-M PFS plots comparing patients with and without STK11 mutation (d) ROC curve and AUC of prediction of 5-year recurrence based on patient TMB values (e) ROC curves of elastic net model built on top-200 highly variable genes in tumor to predict 5-year recurrence (f) ROC curves of elastic net model built on top-200 highly variable genes in normal tissue to predict 5-year recurrence

### Gene expression in tumor-adjacent normal holds significant prognostic information

We then tested whether gene expression obtained from bulk RNA-seq can provide prognostic information by predicting 5-year recurrence. To this end, we constructed an elastic net machine learning model to predict systemic and locoregional recurrence, using nested cross-validation to allow for automatic, unbiased hyper-parameter optimization ensuring no data leakage from training to test sets (see Methods for details). Our analysis determined that models that include transcriptomic information from the normal lung samples outperform models based on tumor histology alone (AUC = 0.83, 95% confidence interval = [0.74-0.92], see **Figure 2e**) and is able to stratify the patients into high- and low-risk groups (PFS log-rank test p-value = 0.007), suggesting that normal lung microenvironment may also contribute to recurrence. The transcriptomic signature in the tumor does not predict recurrence (AUC=0.63, 95% confidence interval = [0.53-0.72]) (**Figure 2f**) and cannot stratify the patients into high- and low-risk groups (PFS log-rank test p-value = 0.456), highlighting the importance of including normal lung samples in our study.

### Co-expressed gene module analysis reveals the activation of inflammatory pathways in tumor-adjacent normal lung tissue

After confirming that somatic mutations do not inform patient outcome and that transcriptomic data from the matched tumor-adjacent normal lung hold significant prognostic information, we set out to characterize the transcriptional programs that are activated in patients that eventually progress. Instead of relying on complex supervised machine learning models (**Figure 2e,f**) with a potentially large number of parameters and questionable capacity to generalize in a clinical setting, we decided to further analyze the 246 matched tumor-normal RNA-seq samples using an unsupervised unbiased approach. Briefly, we selected the top 10,000 most variable genes, scaled their expression across samples and performed dimensionality reduction using UMAP (each point on the UMAP represent a gene, see Methods for details). Unsupervised clustering revealed 20 gene clusters, i.e. co-expressed gene modules, or, simply, modules (**Figure 3a**). We then colored each gene by its log-fold change from normal to tumor samples, revealing clusters of genes with higher expression in the tumor samples (red color) and clusters with higher expression in the normal samples (blue color) as shown in **Figure 3b**. To identify the modules that have overall higher expression in tumor compared to tumor-adjacent normal and *vice versa*, we defined a score for each module as the average scaled gene expression of genes in the module (per patient, per tissue type). Indeed, we found that several modules have significantly higher average expression in the normal samples (modules 1, 2, 5, 6, 7, 8, 9, 11, 17, 19, and 20), while others were more highly expressed in tumor samples (modules 3, 4, 10, 12, 13, 14, 15, 16, and 18) (**Figure 3c**). We then characterized each module based on its association with hallmarks, gene sets with well-defined biological states or processes^21^. The module found to be associated with the highest number of hallmarks was module 20 (**Figure 3d**). Strikingly, although module 20 has a higher score in the normal lung tissue compared to tumor, it was found to be significantly enriched in a large number of hallmarks that are typically linked to cancer, suggesting that tumor-adjacent normal tissue is not entirely normal. In particular, inflammatory signaling pathways (TNF-α, IL-17 and NFκB), IL-2 and IL-6 signaling, interferon-gamma response and hypoxia were found to be highly enriched in module 20 genes. The full list of enriched hallmarks, KEGG pathways and Gene Ontology (GO) terms and all statistics can be found in **Supplementary File 4**.

**Figure 3:**
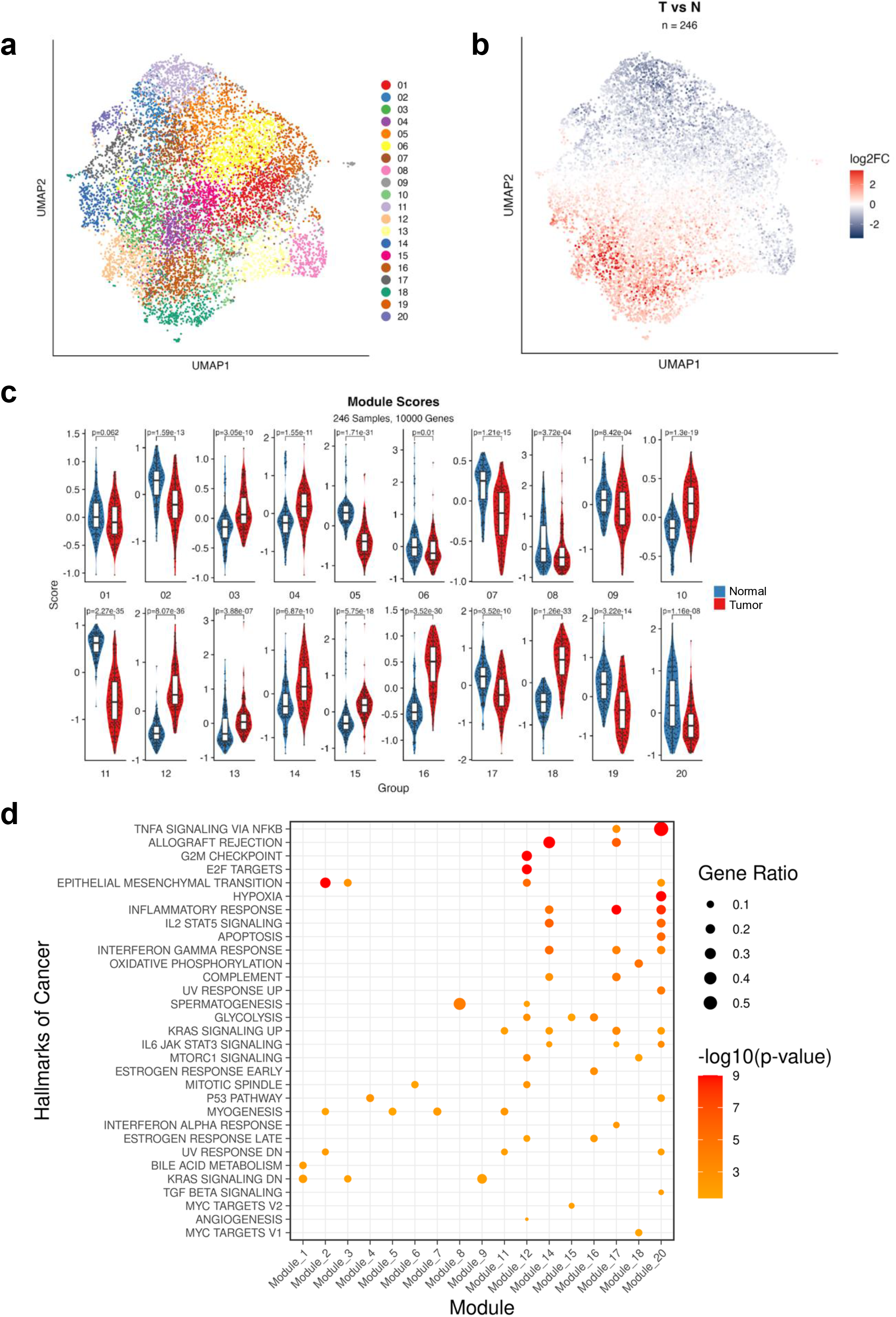
Gene co-expression modules in tumor and normal samples. (a) UMAP representation of 20 gene co-expression modules, (b) UMAP representation annotated by log-fold change tumor vs normal, (c) Boxplots comparing modules scores in tumor and normal samples in each module, (d) Dot plot of enriched KEGG pathways across modules (module 10, 13 and 19 have no highly significant associations).

### Transcriptomic signatures of lung adenocarcinoma progression in tumor and tumor-adjacent normal tissue

Motivated by the observation that inflammatory and other pathways linked to cancer are activated in tumor-adjacent normal, we hypothesized that activation of such pathways and related gene modules, most notably module 20 which was found to be associated with the highest number of cancer-related hallmarks, may inform disease progression. To test this hypothesis, we identified genes that are differentially expressed in the tumor or adjacent normal tissue between the group of patients that eventually progress and the ones that do not. More specifically, patients from our matched tumor-normal cohort were divided into two groups: the progression group comprised all patients with any type disease progression (n=45), while the no progression group comprised all patients that have not progressed with at least 5 years of follow-up time (n=68). Differential expression analysis between the two groups was performed separately on the tumors (**Supplementary Figure 6a**) and the normal lung samples (**Supplementary Figure 6b**). We observed a similar number of significant differentially expressed genes in the two tissue types (672 in tumor and 474 in normal), while the two lists of differentially expressed genes showed minimal overlap, suggesting that the dysregulated pathways in patients that eventually progress are different in the tumor and the tumor-adjacent normal tissue. The results of the differential expression analysis are available as **Supplementary File 5**. We then explored the distribution of differentially expressed genes across the co-expressed gene modules. We colored each gene in the gene module UMAP (**Figure 4a**) by the log-fold change in expression between the progression and no progression groups, separately for the tumor (**Figure 4b**) and the normal samples (**Figure 4c**). Visual inspection and comparison of the UMAPs revealed that upregulated genes in patients that eventually progressed are localized almost exclusively in particular modules, especially in the tumor-adjacent normal lung samples. The most prominent such module is module 20 which has a high percentage of upregulated genes in the normal lung tissue of patients who progress. This is confirmed by module aggregate expression analysis (**Figure 4d, Supplementary Figure 6c**), by calculating the percentages of up- and down-regulated genes across modules in the two tissue types (**Figure 4e**). Clearly, module 20 is highly biased towards upregulated genes in the progressors’ group in the tumor-adjacent normal tissue, but not in the tumor.

**Figure 4:**
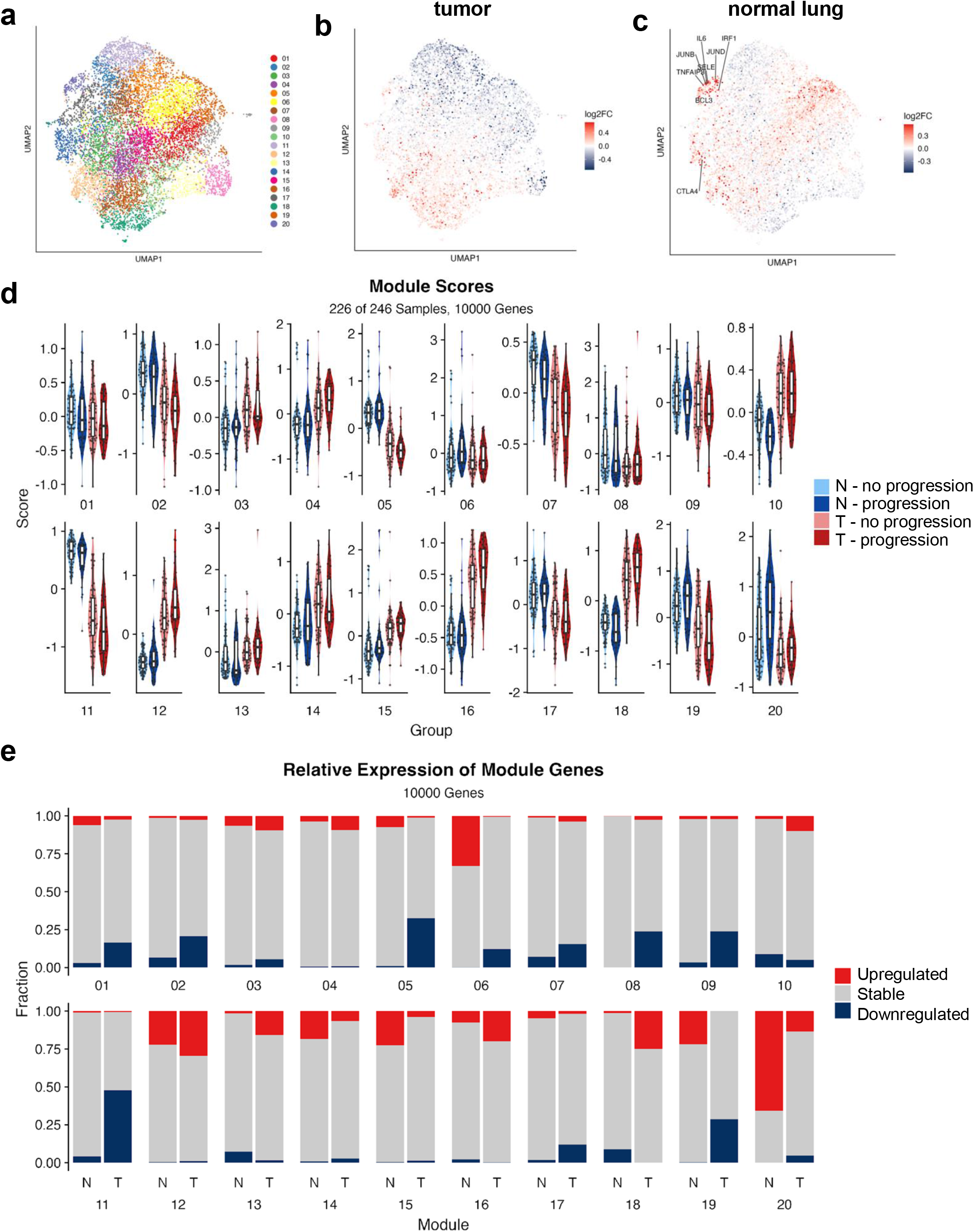
Gene co-expression modules in lung adenocarcinoma progression. (a) UMAP representation of 20 gene co-expression modules, (b) UMAP representation annotated by log-fold change progression (red color) vs no progression (blue color) in tumor samples, (c) UMAP representation annotated by log-fold change progression (red color) vs no progression (blue color) in normal samples, (d) Boxplots comparing modules scores by progression status in tumor (T) and normal (N) tissue in each module, (e) Percentages of up- and down-regulated genes (progression vs no progression) in tumor (T) and normal (N) tissue in each module.

### A multi-modal association map for refined patient classification

Then, focusing first on the tumor-adjacent tissue, we performed a comprehensive association analysis of module scores with demographic, clinical, histologic, genetic and survival data (**Figure 5a**). The only module significantly associated with poor survival is module 20 (**Figure 5b**) and it was found to be an independent predictor of clinical outcome in a multivariate analysis (**Figure 5c**) with a log odds-ratio of 0.725 (p-value = 0.002). Intriguingly, IASLC grade which is part of the updated WHO guidelines of lung adenocarcinoma, was not found significant in the same multivariate analysis. The association map in **Figure 5a** provides a wealth of information that can be used in future bigger studies to not only stratify patients into highly refined groups based on a combination of demographic, clinical, histologic and genetic data, but also generate hypotheses regarding the underlying biological processes and pathways involved by integrating with transcriptomic data from the tumor and the tumor-adjacent normal. For example, modules 7 and 10 are associated with younger patients, are broadly associated with low grade tumors, absence of high-risk histologic patterns (solid and fused grands) and better outcomes. Modules 19 and 20 are associated with older patients and high-grade tumors, although only module 20 was found significantly associated with clinical outcome. Modules 8, 12 and 13 are associated with pleural invasion. Interestingly, none of the modules is associated with mutations, supporting our original hypothesis that the tumor-adjacent normal tissue may be a valuable source of biomarkers for progression, independent of the genetic makeup of the tumor itself. In particular, module 20 activation occurs in patients that progress independent of the driver mutation of their tumors. The same type of association map was generated for the tumor samples (**Supplementary Figure 7a**). Module 20 in the tumor has no significant association with survival in univariate or multivariate analysis (**Supplementary Figure 7b,c**).

**Figure 5:**
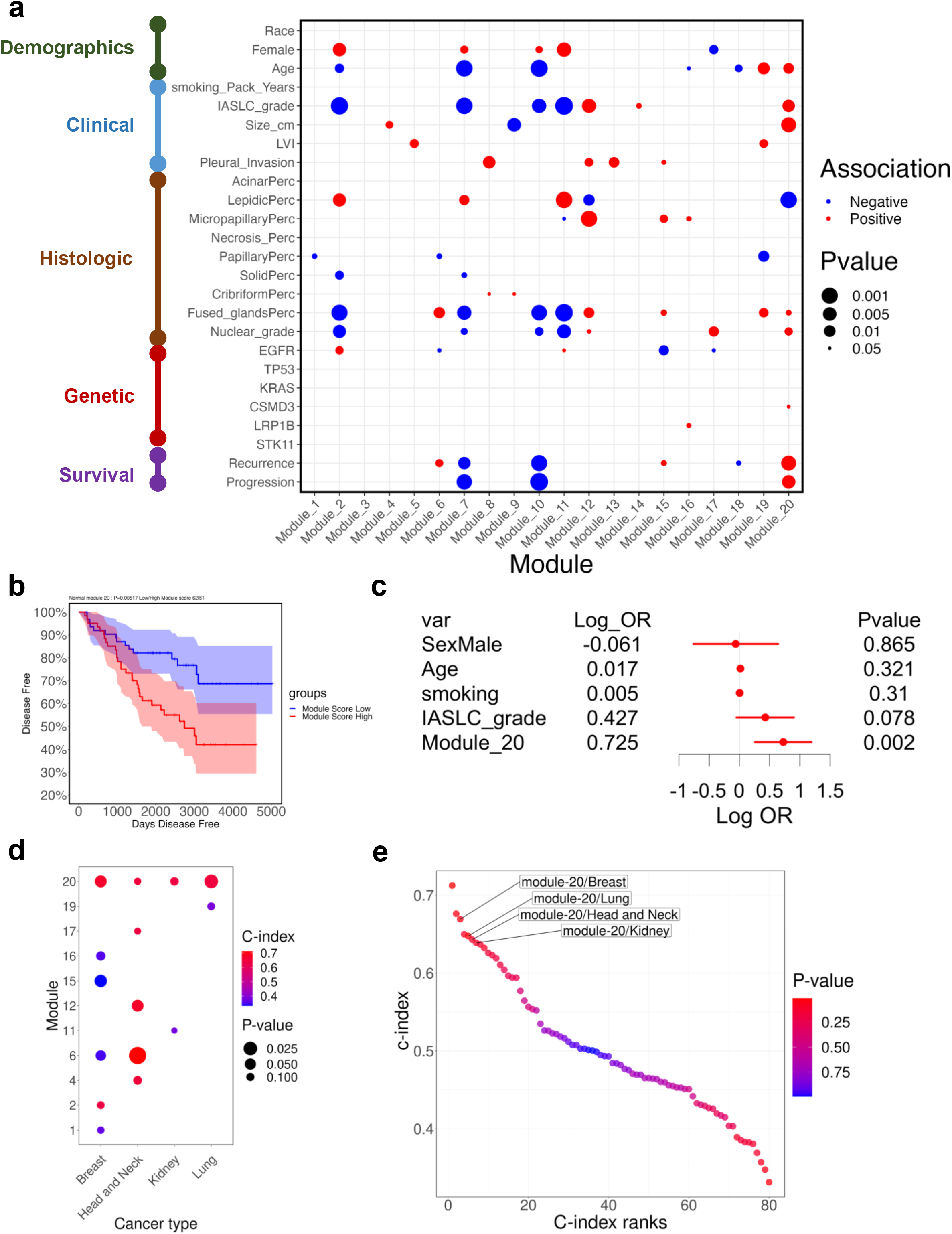
Association of module scores in normal tissue with different variables. (a) Positive and negative associations of demographic, clinical, histologic, genetic and outcomes with module scores in normal tissue (b) K-M PFS curve for patients with high and low module 20 scores in normal tissue (c) Multi-variate modeling of time-to-progression (d) Dot plot of c-index values across modules and TCGA cohorts (e) Elbow plot of c-index values across modules and TCGA cohorts

### Testing the inflammatory module 20 signature on additional cancer types

To further test whether the module 20 inflammatory signature can be more broadly applied to the adjacent normal tissue of other cancer types, we performed an analysis of the data obtained from normal tissue in TCGA. Given the limited number of normal samples in TCGA with RNA-seq data, we were only able to find 4 primary tumor sites with at least 40 tumor-adjacent normal samples across all stages: breast, lung, kidney and head/neck cancer. We calculated c-index values between module scores and progression-free survival for each module and each cancer type (c-index values are higher when high module scores are associated with worse survival). The results of this analysis are shown in **Figure 5d**, demonstrating that module 20 is the only module score that is consistently and significantly associated with poor outcome in all 4 cancer types, as also demonstrated by the elbow plot in **Figure 5e**. Furthermore, Kaplan-Meier curves of high-vs-low module 20 scores, demonstrate that increased module 20 expression is associated with worse survival for both PFS and OS (**Supplementary Figure 8**). Taken together, these findings suggest a prominent role of module 20 in patient progression. As shown above in **Figure 3d**, this module is enriched in inflammatory signaling pathways (TNF-α, IL-17 and NFκB) and hallmarks of cancer (IL-2 and IL-6 signaling, interferon-gamma response and hypoxia), even though it is a module that is more highly expressed in the adjacent normal compared to the actual tumor. This observation suggests that patients who eventually progress, have compromised lungs bearing hallmarks of disease progression that are not necessarily observable in the adjacent tumors.

### Profiling the tumor-adjacent normal tissue at single-cell resolution

In addition to bulk RNA-seq analysis, we utilized single-nucleus RNA-sequencing (snRNA-seq) to analyze the tumor-adjacent normal of our early-stage lung adenocarcinoma matched tumor-normal cohort. We used single-nucleus RNA-seq (snRNA-seq) to profile 23 tumor and 23 matched tumor-adjacent normal samples with left-over material (see Methods for details). Following post-sequencing quality control we were left with 18 tumor and 15 normal snRNA-seq samples (119,629 nuclei). Genotyping analysis of the snRNA-seq data confirmed that these samples match the patient samples used for bulk RNA-seq (**Supplementary Figure 9**). Cells were annotated based on a previous study of lung adenocarcinomas which included normal lung as control^22^ (see Methods). Focusing on the set tumor-adjacent normal samples (51,428 nuclei) (**Supplementary Figure 10a**), we identified the major cell types: epithelial cells, stromal cells, endothelial cells, myeloid cells, T-NK cells, B lymphocytes and MAST cells (**Figure 6a**). The distinct cell lineages were further delineated into more granular subpopulations (**Figure 6b, Supplementary Figure 10b**). Epithelial cells were divided into 5 subtypes: alveolar type 1 and 2 cells (AT1/AT2), club cells and ciliated cells. Stromal cells were divided into 5 subtypes: mesothelial cells, COL13A1 and COL14A1 matrix fibroblasts (FBs), myofibroblasts and pericytes. Endothelial cells (ECs) were divided into 3 subtypes: lymphatic, stalk-like and tip-like ECs. Myeloid cells were divided into 5 subtypes: alveolar macrophages, monocytes, activated dendritic cells (DCs), CD1c DCs and CD207CD1a LCs.

**Figure 6:**
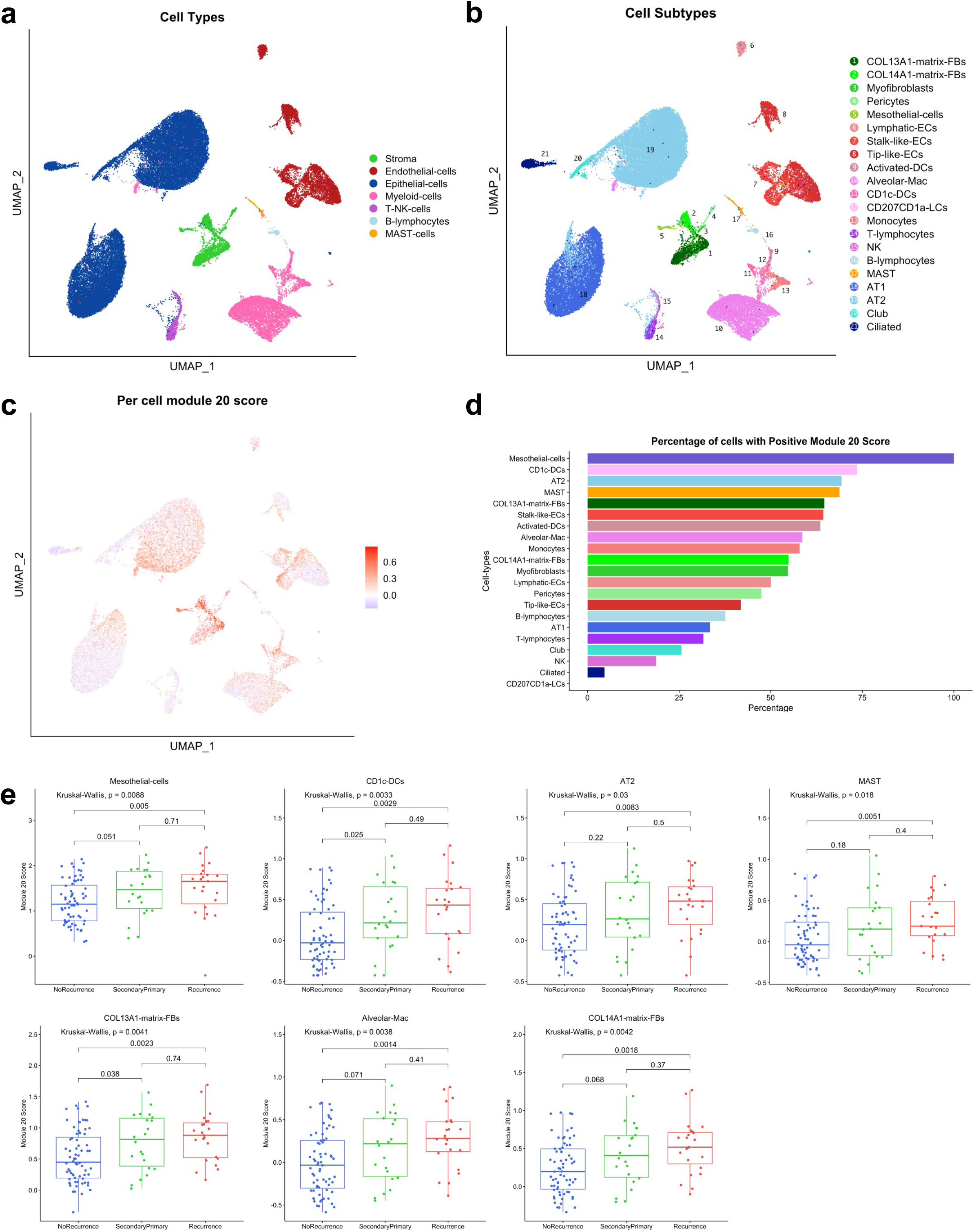
Single-nucleus RNA-seq analysis of adjacent normal. (a) UMAP visualization of all 51,428 adjacent normal nuclei, color-coded based on the broad cell type annotation (b) UMAP visualization of all 51,428 adjacent normal nuclei, color-coded based on the cell subtype annotation (c) UMAP colored by module 20 score (calculated per nucleus) (d) Percentage of cells with a positive module 20 score in each cell subtype (e) Cell subtypes with significantly upregulated expression of the module 20 signature in patients that eventually progress; statistical significance is calculated using the Mann-Whitney test.

### Module 20 is activated in multiple cell types in the tumor-adjacent normal of patients that progress

To test which cell types in the tumor-adjacent normal lung have elevated expression of genes in module 20, we calculated the module 20 score per cell (**Figure 6c**). We observed that the cell type that expressed the highest levels of module 20 genes were mesothelial cells, followed by CD1c DCs, AT2 cells and MAST cells, fibroblasts and alveolar macrophages (**Figure 6d** and **Supplementary Figure 10c**). Recently, mesothelial cells have been shown to form antigen-presenting cancer-associated fibroblasts (apCAFs), which in turn induce naive CD4+ T cells into regulatory T cells in pancreatic cancer^23^. Activation of the module 20 gene signature in AT2 cells (and not in AT1 cells) is also interesting because AT2 cells have been shown to be the cell of origin for lung adenocarcinoma^24^. To further investigate the activation of the module 20 signature in the tumor-adjacent normal tissue of the entire patient cohort, we applied BayesPrism^25^, a Bayesian statistical model that uses single-cell reference to deconvolve bulk RNA-seq expression. Based on our snRNA-seq data, BayesPrism inferred the cell-type composition of our larger bulk RNA-sequencing cohort (**Supplementary Figure 10d**). Overall, we found that the relative abundance of mesothelial cells, and to a lesser extent of monocytes and CD1 DCs, correlated highly with the module 20 score calculated from the bulk RNA-seq data, suggesting an increased production of mesothelial cells in lung tissue with increased overall TNF-α and NFκB signaling (**Supplementary Figure 10e**). We then tested which cell types have upregulated expression of the module 20 signature in patients that progress. For this analysis, we used the inferred gene expression for each cell type in each patient. The results show a concomitant increase, in multiple cell types, of the module 20 score in patients that eventually develop a second primary or recurrence (**Figure 6e**).

## Discussion

Early-stage lung adenocarcinoma is typically treated by surgical resection of the patient’s tumor. While in the majority of cases early intervention can lead to cure, approximately 30% of patients present with disease progression and eventually most of them eventually succumb to metastatic disease. Despite intense efforts to map the genetic landscape of early-stage lung tumors, there has been limited success in discovering accurate biomarkers that can predict progression-free survival. To address this significant unmet need, we proposed that the tumor-adjacent lung of early-stage lung adenocarcinoma patients is an unexplored source of potential biomarkers, and used a unique matched tumor and tumor-adjacent lung adenocarcinoma treatment naive patient cohort to identify molecular signatures associated with progression. To our knowledge, this is the largest such cohort both by size (number of patients) and follow-up time. We profiled both tumor and matched tumor-adjacent specimens using DNA and RNA sequencing and showed that gene expression in tumor-adjacent tissue is the best predictor of disease progression. Tumor heterogeneity across patients is a plausible explanation for this observation: although there are certainly frequently mutated driver genes in lung adenocarcinoma, such as EGFR, KRAS or STK11, only a minority of patients’ tumor is found positive for each of these mutations. Furthermore, co-occurring mutations and the presence of multiple tumor clones and subclones further complicate the already complex mutational landscape. Consequently, there is also a large diversity of transcriptional programs and pathways that are deregulated in tumors. This variability in the tumor transcriptomes is typically supported by PCA plots of tumor and normal samples: tumors are more scattered while normal samples cluster closer together. Therefore, because of the lack of commonly deregulated pathways across tumors, it is not surprising that models of disease progression that are based on tumor only are not accurate. By contrast, we found that tumor-adjacent tissue has a less diverse transcriptional profile, independent of the underlying driver mutations found in the adjacent tumor. This observation leads to the hypothesis that a common set of pathways may be activated in the tumor-adjacent tissue of patients that are at high risk for progression. Indeed, unsupervised discovery of co-expressed gene modules using bulk RNA-sequencing data obtained from our matched cohort uncovered an inflammatory signature (module 20) that can stratify patients into high and low risk groups independent of the underlying mutations found in their tumors. Intriguingly, we demonstrated that the module 20 signature is also associated with poor outcome in several other cancer types, suggesting that a common set of pathways is activated in the tumor-adjacent tissue of tumors that eventually progress. Further supporting our hypothesis, previous studies in other cancers have also suggested that tumor-adjacent tissue has distinct features that could provide prognostic information: hippo-related gene expression in hepatocellular carcinoma^26^, elevated mRNA levels of thymidylate synthase, vascular endothelial growth factor, and EGFR in rectal cancer^27^, different genes being expressed in prostate cancer^28^, and suppression of DMBT1 by cancer cells in squamous cell carcinomas^29^. Another study found that pathways shared among normal tissue adjacent to tumor are altered across different tumor types, and suggests that pro-inflammatory signals from the tumor leads to the stimulation of an inflammatory response in the adjacent endothelium^30^. Moreover, specifically in lung cancer, the concept of “field cancerization” has been explored by Spira et al with their investigations which demonstrate the utility of transcriptomic profiles from proximal airways as an adjunct to routine bronchoscopy for the diagnosis of the indeterminate pulmonary nodule^20^.

Finally, analysis of the transcriptome of tumor-adjacent lung at single-cell resolution revealed that these inflammatory pathways are activated in specific cell types, mostly mesothelial cells, followed by CD1 dendritic cells, alveolar type 2 cells and MAST cells. Two major pathways were identified as highly enriched in the module 20 signature: (1) the TNF-α pathway with genes IL6; JUNB, IRF1, SELE and BCL3 overexpressed in patients who eventually progressed, and, (2) the IL17 pathway with genes JUND, TNFAIP3 and IL6 overexpressed. These two pathways suggest the provocative idea that patients with early-stage lung cancer may benefit from neoadjuvant therapy after their tumors are resected, such as TNF-α blockers or IL-17 inhibitors. Alternatively, the activation of inflammatory pathways in tumor-adjacent tissue may indicate that micrometastases have already occurred at undetectable levels. In such a scenario, there is evidence that blocking inflammation can help eradicate micrometastasis^31^. In addition, early-stage lung adenocarcinoma patients at high risk for progression may benefit from immunotherapy. In conclusion, our studies suggest that molecular profiling of tumor-adjacent tissue can identify patients that are at high risk for progression and may help indicate appropriate neoadjuvant therapies for patients at risk.

## Acknowledgements

We would like to thank the Genome Technology Center (GTC) for expert library preparation and sequencing, the Applied Bioinformatics Laboratories (ABL) for providing bioinformatics support and the Center for Biospecimen Research and Development (CBRD) Histology Core Facility (RRID:SCR_018304). GTC, ABL and CBRD are shared resources partially supported by the Cancer Center Support Grant P30CA016087 at the Laura and Isaac Perlmutter Cancer Center. This work has used computing resources at the NYU School of Medicine High Performance Computing (HPC) Facility. LNS is supported by R37 CA244775 (LNS, NCI/NIH), PACT grant (LNS, FNIH), American Association for Cancer Research Grant (HP/LNS). HIP is supported by NCI/NIH Early Detection Research Network Grant 1U01CA214195. This research was also supported by Roche Access to Distinguished Scientists (ROADS) Programme.

## Methods

### Specimen collection

Snap frozen Stage I lung cancer tumor and matching adjacent lung specimens (within the same lobe, segment, or wedge resection) from 143 patients having R0 resection with lymph node dissection were prospectively collected and archived at −80°C from 2005-2015 under an IRB approved NYULH protocol (i8896). Subjects who received antibiotics (except peri-operative), steroids, radiation, immunotherapy, or chemotherapy within the month prior to surgery were excluded from the study. Subjects were assessed postoperatively with an in-person clinic visit and surveillance chest CT every three months after surgery for two years, every six months for the third year and then yearly.

### Histologic characterization

Histological sections of the pulmonary adenocarcinomas were evaluated in formalin fixed paraffin embedded tissue. The percentage of each histological growth pattern (lepidic, acinar, papillary, solid, micropapillary, and complex glandular patterns (cribriform and fused glands) were recorded in 5% increment for each tumor to a sum of 100% as suggested by the current WHO classification of lung tumor^32^.

### DNA sequencing

DNA sequencing of lung tumors, adjacent normal lung samples and matched normal DNA extracted from blood, was performed using CLIA certified, clinically validated NYU Genome PACT assay for analysis of mutations and copy number changes. NYU Genome PACT is NYS approved, FDA-cleared custom-built, hybrid capture NGS assay analyzing all exons of 607 genes and TERT promoter, using IDT probes, sequenced on Illumina NextSeq 550 system (Illumina, San Diego, CA), with starting DNA input 200 ng, and average depth of sequencing 300x.

### DNA sequencing analysis

Sequencing results were demultiplexed and converted to FASTQ format using Illumina bcl2fastq software. The FASTQ files were processed using Seq-N-Slide pipeline^33^. The reads were adapter and quality trimmed with Trimmomatic^34^ and then aligned to the human reference genome (build hg38/GRCh38) using the Burrows-Wheeler Aligner with the BWA-MEM algorithm^35^. Low confidence mappings (mapping quality <10) and duplicate reads were removed using Sambamba^36^. Further local indel realignment and base-quality score recalibration were performed using the Genome Analysis Toolkit (GATK)^37^. Somatic variants in matched samples were called with Mutect^38^ and Strelka^39^. ANNOVAR^40^ was used to annotate variants with genomic context such as functional consequence on genes and identify presence in public variant databases. The mean depth of coverage across all samples was 935X. Variant calls required >1% VAF, a minimum of 100 total reads, 5 alt reads, and a VAF >5 times that of a matched normal blood. To further reduce the likelihood of false positives, only known somatic variants present in Catalogue Of Somatic Mutations in Cancer (COSMIC)^41^ v94 and with a population frequency of <0.1% based on gnomAD^42^ v2.1.1 were retained.

### RNA sequencing

The quantity and quality of total RNA was assessed on a 2100 BioAnalyzer instrument (Agilent Technologies, Inc.). 1 ng of total RNA was used to prepare libraries using Trio RNA-Seq library prep kit (Tecan Genomics, Inc., part number 0506-96, mammalian rRNA Deplete) following the manufacturer’s instructions. Briefly, the library prep consists of the following steps: DNase treatment to remove genomic DNA, first strand and second strand cDNA synthesis from the input RNA, single primer isothermal amplification (SPIA) of the resultant cDNAs, enzymatic fragmentation and construction of unique barcoded libraries, PCR library amplification and a final step to remove rRNA transcripts. The Agencourt AMPure XP bead (Beckman Coulter) purified libraries were quantified using qPCR and the size distribution was checked using Agilent TapeStation 2200. The libraries were pooled and run on an Illumina S4 flow cell on a NovaSeq as paired end 100.

### RNA sequencing analysis

Sequencing results were demultiplexed and converted to FASTQ format using Illumina bcl2fastq software. The FASTQ files were processed using Seq-N-Slide pipeline^33^. The sequencing reads were adapter and quality trimmed with Trimmomatic^34^ and then aligned to the human reference genome (build hg38/GRCh38) using the splice-aware STAR aligner^43^. The featureCounts program^43^ was utilized to generate counts for each gene based on how many aligned reads overlap its exons. Useable samples were defined as those with more than 30% uniquely mapped reads, less than 50% of bases aligned to rRNA sequences, and more than 5 million assigned counts. The counts were then normalized and used to test for differential expression using negative binomial generalized linear models implemented by the DESeq2 R package^45^.

### Clustering patient samples by genotype

Sample relatedness to ensure that data from the same patients was correctly labeled was computed using Somalier^46^, which analyzes ancestry based on common variants across all human populations and calculates pairwise coefficient of relationship. Sample pairs with less than 50 sites were excluded. Pairwise relatedness values less than 0 were set to 0 for visualization and hierarchical clustering.

### Machine learning classifier for 5-year recurrence

We trained a logistic regression model with elastic net penalty to classify patients that recur vs those that do not based on their gene expression. Our machine learning method combines hard filtering (200 most variable genes) with soft filtering (elastic net regression) and therefore we utilized a nested cross-validation scheme to get an unbiased estimate of its performance and avoid data leakage. At a high-level we use the following steps:

1. For a given outer train-test 10-fold split:

a. we selected top-N genes based on the training data (details below)
b. we used an inner 10-fold on the training data to optimize the parameters (details below)
c. the optimal model, as determined by inner cross-validation, was then applied to the test data of that split
2. Then, we combined the predictions of the individual test sets across all splits (thus covering the entire cohort), and we used these predictions to:

a. generate ROC curves and calculate AUCs
b. split patients into high-low risk based on the recurrence prediction and generate Kaplan-Meier plots.

More specifically, the outer CV-split defines the X matrix while the inner CV fits the β, λ, α parameters. In more detail, in the outer loop we identify the 200 most variable (according to the median absolute deviation) genes of the training split and use them to fit a logistic regression model elastic net regularization. The fitting of the model takes place in the inner (10-fold) cross-validation using 21 potential values for α (a_n=(n/20)^2 for n = 0, …, 20) while the λ values were automatically adjusted by the glmnet package. The best model, in terms of mean cross-validated error, from the inner CV is then used to classify the test cases of the outer split. Finally, the predicted probability of recurrence for all the test sets were combined and used to estimate the ROC curves and plot the Kaplan-Meier curves. The formula for the logistic regression with elastic net penalty is shown below:

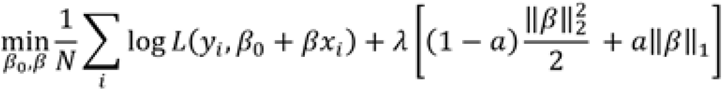

Where: L(y,y^) is the likelihood of the binomial distribution and β_0,β are the parameters to be tuned, x_i is the expression vector for patient i whose dimensions are the 200 most variable genes across all patients, λ controls the regularization penalty, α the trade-off between lasso and ridge regression.

### Gene co-expression analysis

The expression counts were transformed using variance stabilizing transformation (VST) implemented in DESeq2. Only protein-coding genes as identified by GENCODE were retained. Genes were further subset to the 10,000 most variable. Principal components analysis (PCA) was performed on a data matrix of values that were scaled and centered for each gene. The first 10 PCs were used for clustering and UMAP visualization. Gene modules were determined using Partitioning Around Medoids (PAM) clustering implemented in cluster R package with a k=20 and pamonce=5. UMAP was generated using the uwot R package with n_neighbors=10 and min_dist=0.3. The module scores were defined as the average of the z-scores of genes within each module.

### Association of module scores with demographic, clinical, histologic, genetic and outcome variables

For the 20 module scores for all patients, we computed the correlation of them with different demographic, histological, clinical and mutational status features. Then the correlation significance level of each module vs. other features was plotted as a dot plot using R package ggplot2 (version 3.3.6).

### Nucleus isolation and sequencing

Nuclei were prepared for 10× Genomics-based single nuclei RNA seq analysis according to a previously published protocol^47^. Briefly, each frozen sample was thawed and macerated in CST buffer for 10 minutes, filtered (70 micron pluriStrainer) and spun at 500g for 5 min at 4°C to pellet nuclei. Nuclei were resuspended in the same buffer without detergent, filtered (10 micron pluriStrainer) and counted using AOPI on a Nexcelom Cellometer. Approximately 10,000 nuclei were loaded immediately into each channel of a 10x Chromium chip (10× Genomics) using 5-prime v1.1 chemistry according to the manufacturer’s protocol (10× Genomics #CG000208). The resulting cDNA and indexed libraries were checked for quality on an Agilent 4200 TapeStation and then quantified and pooled for sequencing on an Illumina NovaSeq 6000.

### snRNA-seq data preprocessing

Sequencing reads were trimmed of adapter sequences using cutadapt^48^. Barcode processing and gene quantification was performed with STARsolo 2.7.3a^49^ using the GRCh38 human reference transcriptome (refdata-cellranger-GRCh38-3.0.0 provided by 10x Genomics). STARsolo pre-mRNA counts were used to generate the gene-barcode matrix. Further analysis including the identification of highly variable genes, dimensionality reduction, standard unsupervised clustering algorithms, and the discovery of differentially expressed genes was performed using Seurat 4.0^50^ and streamlined as an R package (available at https://github.com/igordot/scooter).

Nuclei were filtered to only include those with >500 detectable genes, >1000 UMIs, and <10% of transcripts coming from mitochondrial genes. The UMI counts were normalized by the total number of UMIs per nucleus, multiplied by a scale factor of 10,000, and log-transformed. Likely doublets/multiplets were identified and removed using the scDblFinder package^51^.

### Dimensionality reduction and annotation

To visualize the data, the dimensionality of the scaled integrated data matrix was further reduced to project the nuclei in two-dimensional space using PCA followed by uniform manifold approximation and projection (UMAP)^52^ using top 50 PCs and 30 nearest neighbors to define the local neighborhood size with a minimum distance of 0.3. The resulting PCs were also used as a basis for partitioning the dataset into clusters using a smart local moving (SLM) community detection algorithm^53^. A range of resolutions (0.1-10) was utilized to establish a sufficient number of clusters.

Nuclei were annotated using a previous study of lung adenocarcinomas as a reference^22^. Brain metastasis samples were removed from the reference dataset. SingleR^54^ annotation was performed on the aggregated cluster profiles using 86 clusters (resolution of 3) with cell type and cell subtype labels.

Further analysis was performed on 15 normal samples (51,428 nuclei). To account for biological and technical batch differences between individual patients and scRNA-seq libraries, the Seurat anchor-based integration method for merging datasets that identify pairwise correspondences between cell pairs across datasets to transform them into a shared space was utilized. The 2,000 most variable genes based on standardized variance were selected for canonical correlation analysis (CCA) as an initial dimensional reduction. The integration anchors were then identified based on the first 30 dimensions and used to generate a new dimensional reduction for further analysis. Seurat’s AddModuleScore function was used to quantify gene set expression in each nucleus.

### Bulk RNA-seq deconvolution

BayesPrism (version 2.0) was used for deconvolution of the normal bulk RNA samples^25^. Mitochondrial and ribosomal protein coding genes were excluded from deconvolution analysis. To increase the signal-to-noise ratio, we also removed lowly transcribed genes, leaving us with 19,816 genes. To reduce batch effects and decrease computational time, we retained only protein coding genes for a total gene count of 13,972 for the single-nuclei reference. Next, cells were labeled according to the cell subtypes as identified by a previous group^22^. For BayesPrism, the parameter key was set to NULL to indicate there were no malignant cells in the reference and all 21 cell types are treated equally. Final Gibbs theta values were used to estimate the fraction of each cell type. We extracted the posterior mean of each cell-type specific gene expression for the outputted count matrix, *Z* for every cell type. Next, we computed the z-score for all genes across our cell types of interest. The module 20 score was then defined as the average of z-scores of the module genes. A Mann-Whitney test was run between the progression (second primary or recurrence) and no progression groups for each cell-type. P-values<0.01 were considered statistically significant.

## Supplementary Files

Supplementary File 1: Cohort characteristics

Supplementary File 2: DNA sequencing quality assessment

Supplementary File 3: RNA sequencing quality assessment

Supplementary File 4: Module gene set enrichment analysis (KEGG pathways, Gene Ontology and HALLMARKS)

Supplementary File 5: Differential gene expression analysis

## References

1. Wu CF, Fu JY, Yeh CJ, et al. Recurrence Risk Factors Analysis for Stage I Non-small Cell Lung Cancer. Medicine (Baltimore). 2015;94(32):e1337.

2. Moreira AL, Ocampo PS, Xia Y, et al. A grading system for invasive pulmonary adenocarcinoma: a proposal from the International Association for the Study of Lung Cancer Pathology Committee. Journal of Thoracic Oncology. 2020;15(10):1599–1610.

3. Luo J, Wang R, Han B, et al. Solid predominant histologic subtype and early recurrence predict poor postrecurrence survival in patients with stage I lung adenocarcinoma. Oncotarget. 2017;8(4):7050–7058.

4. Wang X, Janowczyk A, Zhou Y, et al. Prediction of recurrence in early stage non-small cell lung cancer using computer extracted nuclear features from digital H&E images. Sci Rep. 2017;7(1):13543.

5. Yu K-H, Zhang C, Berry GJ, et al. Predicting non-small cell lung cancer prognosis by fully automated microscopic pathology image features. Nature Communications. 2016;7(1):12474.

6. Jones GD, Brandt WS, Shen R, et al. A genomic-pathologic annotated risk model to predict recurrence in early-stage lung adenocarcinoma. JAMA surgery. 2021;156(2):e205601–e205601.

7. Cho SH, Yoon S, Lee DH, et al. Recurrence-associated gene signature in patients with stage I non-small-cell lung cancer. Scientific Reports. 2021;11(1):19596.

8. He Q, Xin P, Zhang M, et al. The impact of epidermal growth factor receptor mutations on the prognosis of resected non-small cell lung cancer: a meta-analysis of literatures. Transl Lung Cancer Res. 2019;8(2):124–134.

9. Lu Y, Wang L, Liu P, et al. Gene-expression signature predicts postoperative recurrence in stage I non-small cell lung cancer patients. PLoS One. 2012;7(1):e30880.

10. Liljedahl H, Karlsson A, Oskarsdottir GN, et al. A gene expression-based single sample predictor of lung adenocarcinoma molecular subtype and prognosis. Int J Cancer. 2021;148(1):238–251.

11. Fahrmann JF, Grapov D, Phinney BS, et al. Proteomic profiling of lung adenocarcinoma indicates heightened DNA repair, antioxidant mechanisms and identifies LASP1 as a potential negative predictor of survival. Clinical Proteomics. 2016;13(1):31.

12. Chen G, Gharib TG, Wang H, et al. Protein profiles associated with survival in lung adenocarcinoma. Proc Natl Acad Sci U S A. 2003;100(23):13537–13542.

13. Billatos E, Vick JL, Lenburg ME, et al. The Airway Transcriptome as a Biomarker for Early Lung Cancer Detection. Clin Cancer Res. 2018;24(13):2984–2992.

14. Slaughter DP, Southwick HW, Smejkal W. Field cancerization in oral stratified squamous epithelium; clinical implications of multicentric origin. Cancer. 1953;6(5):963–968.

15. Blomquist T, Crawford EL, Mullins D, et al. Pattern of antioxidant and DNA repair gene expression in normal airway epithelium associated with lung cancer diagnosis. Cancer Res. 2009;69(22):8629–8635.

16. Spira A, Beane JE, Shah V, et al. Airway epithelial gene expression in the diagnostic evaluation of smokers with suspect lung cancer. Nat Med. 2007;13(3):361–366.

17. Franklin WA, Gazdar AF, Haney J, et al. Widely dispersed p53 mutation in respiratory epithelium. A novel mechanism for field carcinogenesis. J Clin Invest. 1997;100(8):2133–2137.

18. Tang X, Shigematsu H, Bekele BN, et al. EGFR tyrosine kinase domain mutations are detected in histologically normal respiratory epithelium in lung cancer patients. Cancer Res. 2005;65(17):7568–7572.

19. Kadara H, Fujimoto J, Yoo SY, et al. Transcriptomic architecture of the adjacent airway field cancerization in non-small cell lung cancer. J Natl Cancer Inst. 2014;106(3):dju004.

20. Silvestri GA, Vachani A, Whitney D, et al. A Bronchial Genomic Classifier for the Diagnostic Evaluation of Lung Cancer. N Engl J Med. 2015;373(3):243–251.

21. Liberzon A, Birger C, Thorvaldsdóttir H, et al. The Molecular Signatures Database (MSigDB) hallmark gene set collection. Cell Syst. 2015;1(6):417–425.

22. Kim N Kim HK, Lee K, et al. Single-cell RNA sequencing demonstrates the molecular and cellular reprogramming of metastatic lung adenocarcinoma. Nature Communications. 2020;11(1):2285.

23. Huang H, Wang Z, Zhang Y, et al. Mesothelial cell-derived antigen-presenting cancer-associated fibroblasts induce expansion of regulatory T cells in pancreatic cancer. Cancer Cell. 2022;40(6):656–673.e657.

24. Sainz de Aja J, Dost AFM, Kim CF. Alveolar progenitor cells and the origin of lung cancer. J Intern Med. 2021;289(5):629–635.

25. Chu T, Wang Z, Pe’er D, et al. Cell type and gene expression deconvolution with BayesPrism enables Bayesian integrative analysis across bulk and single-cell RNA sequencing in oncology. Nat Cancer. 2022;3(4):505–517.

26. Pan Q, Qin F, Yuan H, et al. Normal tissue adjacent to tumor expression profile analysis developed and validated a prognostic model based on Hippo-related genes in hepatocellular carcinoma. Cancer Med. 2021;10(9):3139–3152.

27. Schneider S, Park DJ, Yang D, et al. Gene expression in tumor-adjacent normal tissue is associated with recurrence in patients with rectal cancer treated with adjuvant chemoradiation. Pharmacogenet Genomics. 2006;16(8):555–563.

28. Zhou R, Feng Y, Ye J, et al. Prediction of Biochemical Recurrence-Free Survival of Prostate Cancer Patients Leveraging Multiple Gene Expression Profiles in Tumor Microenvironment. Frontiers in Oncology. 2021;11.

29. Singh P, Banerjee R, Piao S, et al. Squamous cell carcinoma subverts adjacent histologically normal epithelium to promote lateral invasion. J Exp Med. 2021;218(6).

30. Aran D, Camarda R, Odegaard J, et al. Comprehensive analysis of normal adjacent to tumor transcriptomes. Nature Communications. 2017;8(1):1077.

31. Panigrahy D, Gartung A, Yang J, et al. Preoperative stimulation of resolution and inflammation blockade eradicates micrometastases. J Clin Invest. 2019;129(7):2964–2979.

32. Nicholson AG, Tsao MS, Beasley MB, et al. The 2021 WHO Classification of Lung Tumors: Impact of Advances Since 2015. J Thorac Oncol. 2022;17(3):362–387.

33. Dolgalev I. Seq-N-Slide (v22.01). Zenodo 2022.

34. Bolger AM, Lohse M, Usadel B. Trimmomatic: a flexible trimmer for Illumina sequence data. Bioinformatics. 2014;30(15):2114–2120.

35. Li H, Durbin R. Fast and accurate short read alignment with Burrows-Wheeler transform. Bioinformatics. 2009;25(14):1754–1760.

36. Tarasov A, Vilella AJ, Cuppen E, et al. Sambamba: fast processing of NGS alignment formats. Bioinformatics. 2015;31(12):2032–2034.

37. McKenna A, Hanna M, Banks E, et al. The Genome Analysis Toolkit: a MapReduce framework for analyzing next-generation DNA sequencing data. Genome Res. 2010;20(9):1297–1303.

38. Cibulskis K, Lawrence MS, Carter SL, et al. Sensitive detection of somatic point mutations in impure and heterogeneous cancer samples. Nat Biotechnol. 2013;31(3):213–219.

39. Kim S, Scheffler K, Halpern AL, et al. Strelka2: fast and accurate calling of germline and somatic variants. Nat Methods. 2018;15(8):591–594.

40. Wang K, Li M, Hakonarson H. ANNOVAR: functional annotation of genetic variants from high-throughput sequencing data. Nucleic Acids Res. 2010;38(16):e164.

41. Tate JG, Bamford S, Jubb HC, et al. COSMIC: the Catalogue Of Somatic Mutations In Cancer. Nucleic Acids Research. 2018;47(D1):D941–D947.

42. Karczewski KJ, Francioli LC, Tiao G, et al. The mutational constraint spectrum quantified from variation in 141,456 humans. Nature. 2020;581(7809):434–443.

43. Dobin A, Davis CA, Schlesinger F, et al. STAR: ultrafast universal RNA-seq aligner. Bioinformatics. 2013;29(1):15–21.

44. Liao Y, Smyth GK, Shi W. featureCounts: an efficient general purpose program for assigning sequence reads to genomic features. Bioinformatics. 2014;30(7):923–930.

45. Love MI, Huber W, Anders S. Moderated estimation of fold change and dispersion for RNA-seq data with DESeq2. Genome Biol. 2014;15(12):550.

46. Pedersen BS, Bhetariya PJ, Brown J, et al. Somalier: rapid relatedness estimation for cancer and germline studies using efficient genome sketches. Genome Med. 2020;12(1):62.

47. Drokhlyansky E, Smillie CS, Van Wittenberghe N, et al. The Human and Mouse Enteric Nervous System at Single-Cell Resolution. Cell. 2020;182(6):1606-1622.e1623.

48. Martin M. Cutadapt removes adapter sequences from high-throughput sequencing reads. 2011. 2011;17(1):3.

49. Kaminow B, Yunusov D, Dobin A. STARsolo: accurate, fast and versatile mapping/quantification of single-cell and single-nucleus RNA-seq data. bioRxiv. 2021:2021.2005.2005.442755.

50. Hao Y, Hao S, Andersen-Nissen E, et al. Integrated analysis of multimodal single-cell data. Cell. 2021;184(13):3573–3587.e3529.

51. Germain PL, Lun A, Garcia Meixide C, et al. Doublet identification in single-cell sequencing data using scDblFinder. F1000Res. 2021;10:979.

52. Becht E, McInnes L, Healy J, et al. Dimensionality reduction for visualizing single-cell data using UMAP. Nat Biotechnol. 2018.

53. Waltman L, van Eck NJ. A smart local moving algorithm for large-scale modularity-based community detection. The European Physical Journal B. 2013;86(11):471.

54. Aran D, Looney AP, Liu L, et al. Reference-based analysis of lung single-cell sequencing reveals a transitional profibrotic macrophage. Nature Immunology. 2019;20(2):163–172.

