## Supplemental Figures for "Inflammation in the tumor-adjacent lung as a predictor of clinical outcome in lung adenocarcinoma": FIGURES SUPP.pdf

### Supplementary Figure 1

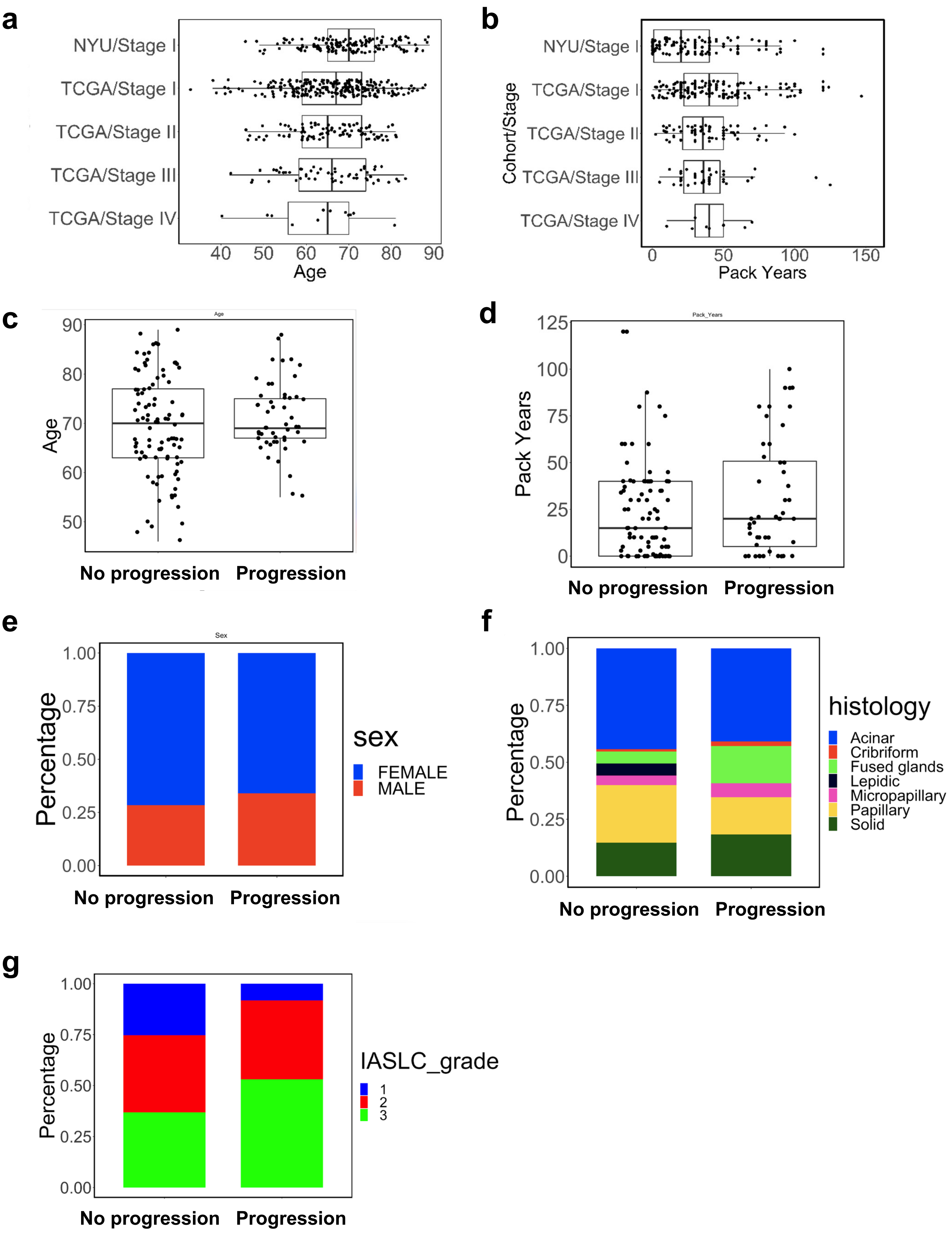

**Supplementary Figure 1. Additional cohort characteristics.**

(a) Patient age distributions represented as boxplots in NYU and TCGA cohorts (by stage),
(b) Patient pack year distributions represented as boxplots in NYU and TCGA cohorts (by stage),
(c) Patient age by progression status represented as boxplots (NYU cohort),
(d) Patient pack year distributions by progression status represented as boxplots (NYU cohort only),
(e) Percentage of male and female patients by progression status (NYU cohort),
(f) Breakdown of histologic types by progression status (NYU cohort),
(g) Tumor grade by progression status (NYU cohort),

#### Supplementary Figure 2

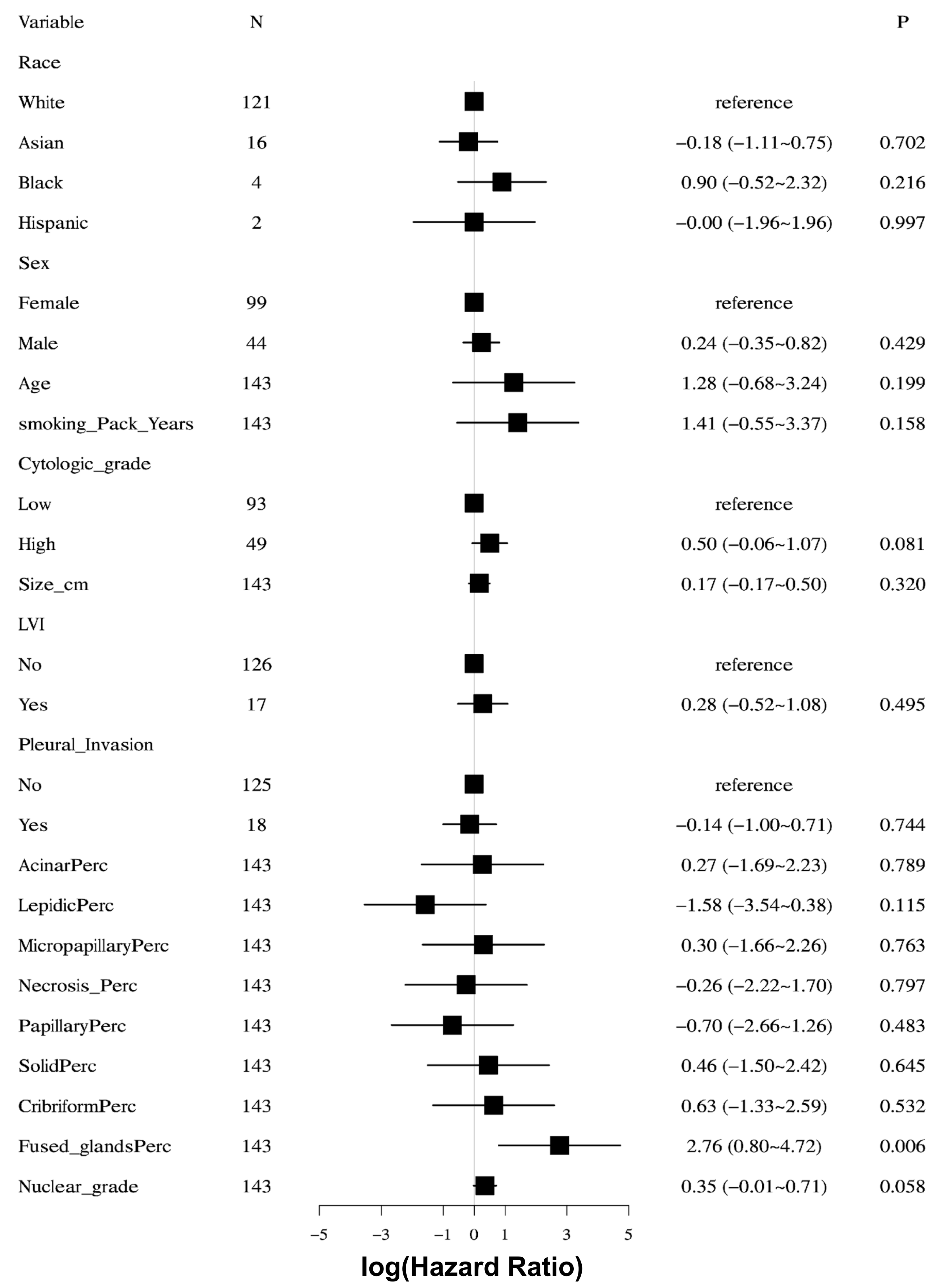

**Supplementary Figure 2. Cox regression on clinicodemographic variables.**

#### Supplementary Figure 3

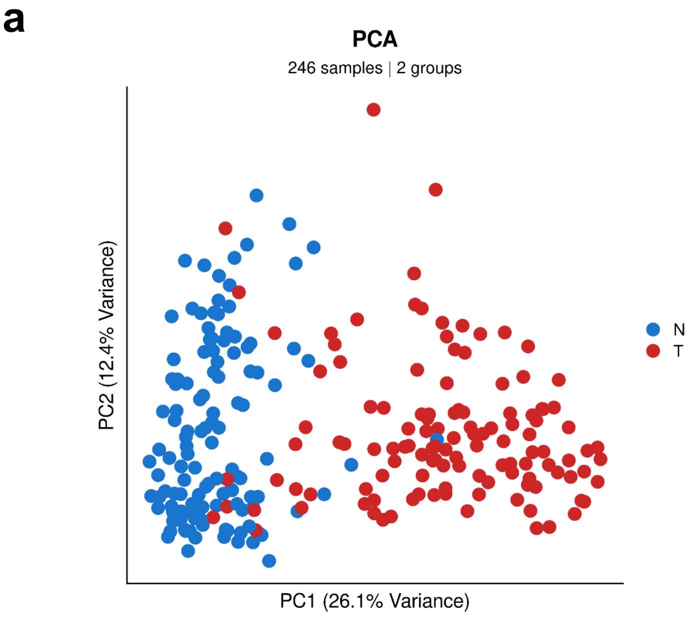


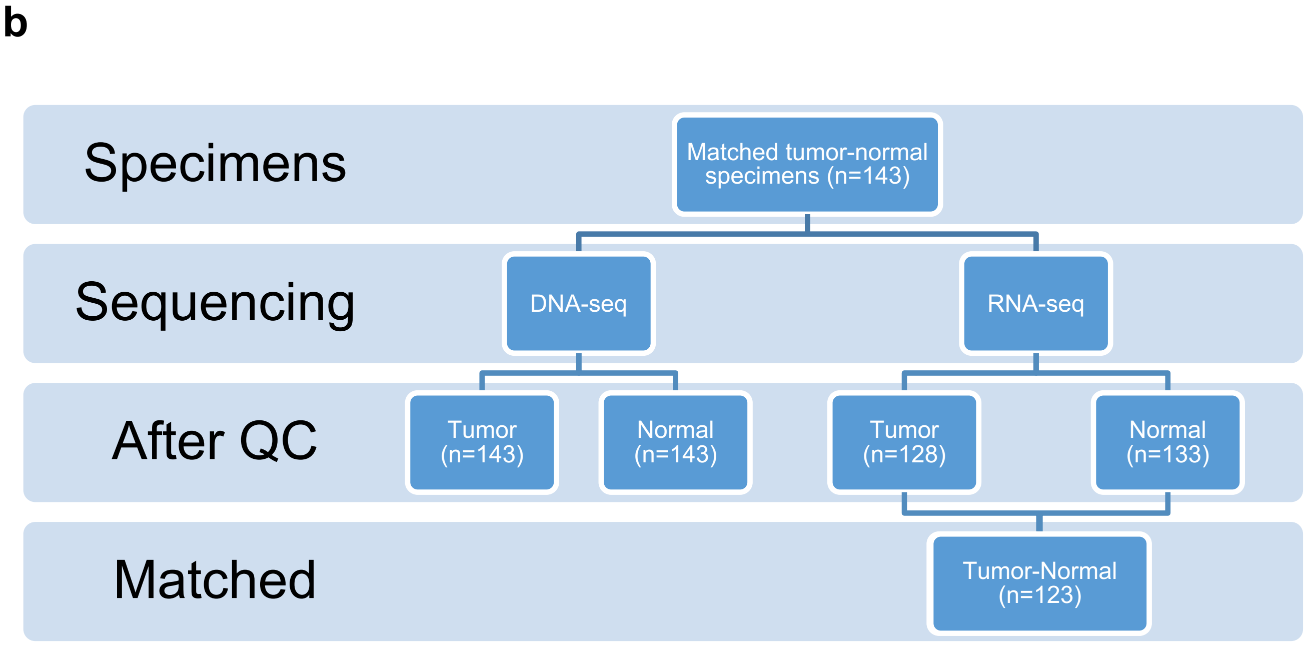

**Supplementary Figure 3. Sequencing and quality control workflow.**

(a) PCA of tumor-normal RNA samples

(b) Number of normal and tumor samples with DNA-seq and RNA-seq

### Supplementary Figure 4

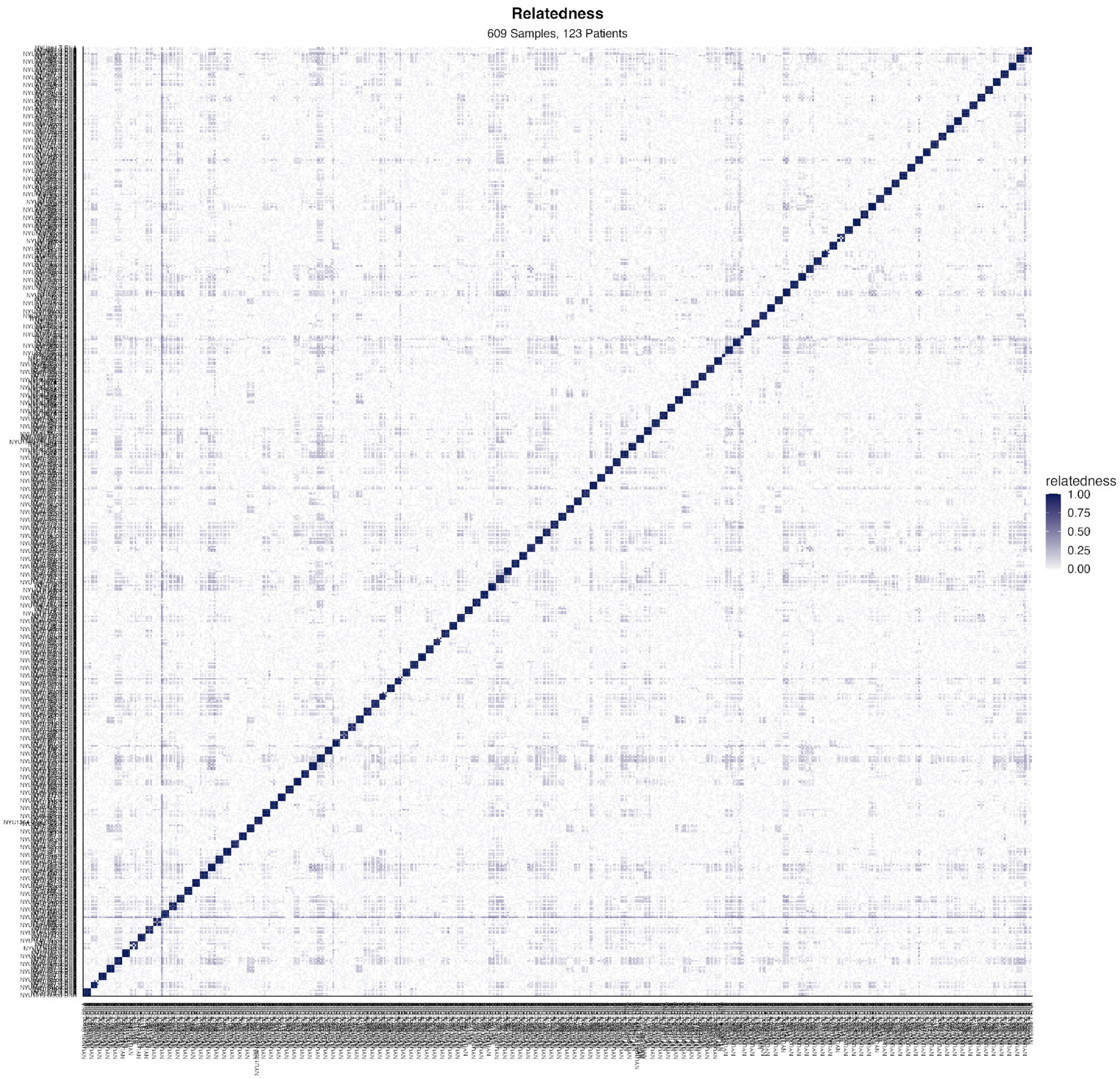

**Supplementary Figure 4. Clustering of all patient samples by genotype.** Pairwise similarity of variants called on all available samples. Each patient has up to 5 sequenced samples: DNA-seq of tumor, normal and blood samples, and RNA-seq of tumor, normal samples. Samples from the same patient cluster together as expected.

### Supplementary Figure 5

**a**

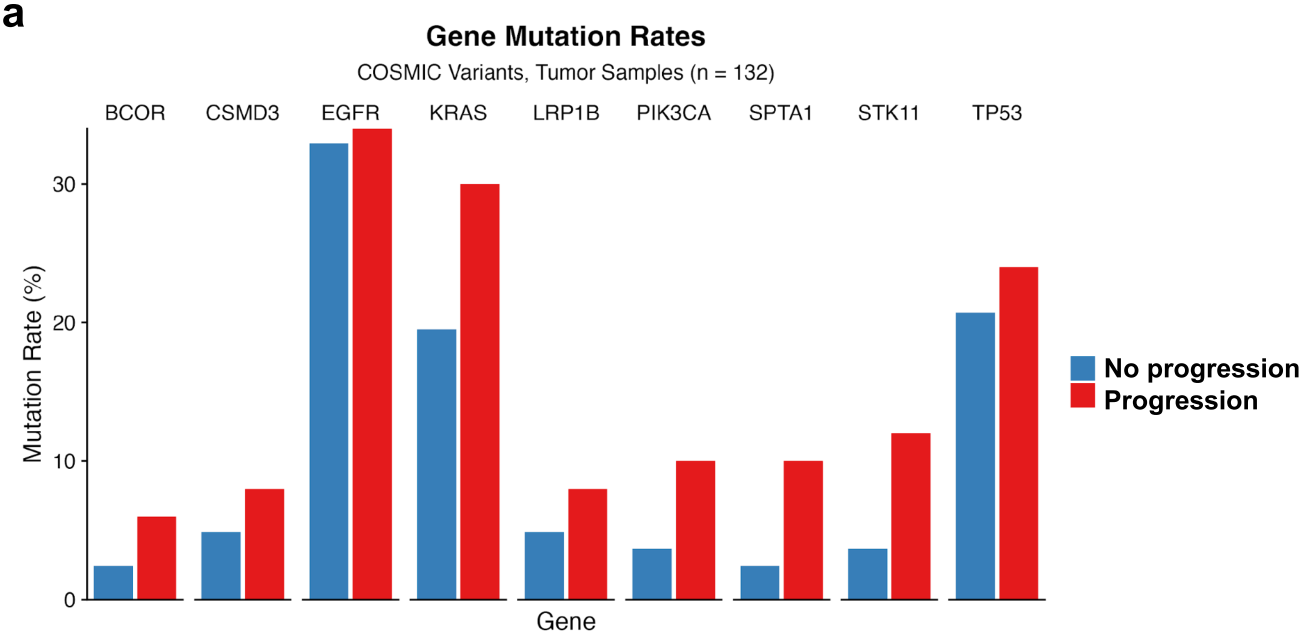


**b**

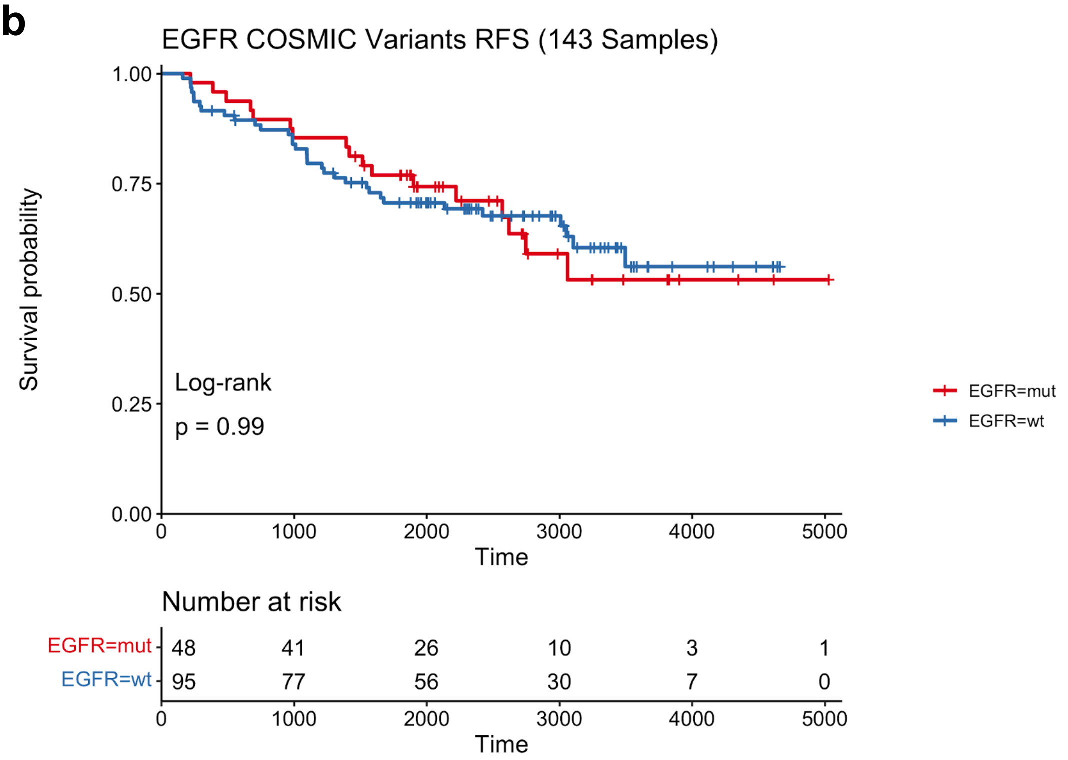

**Supplementary Figure 5: Mutations and disease progression.**

(a) Gene mutation rates by progression status.

(b) Kaplan-Meier progression-free survival (PFS) plots comparing patients with and without EGFR mutation

### Supplementary Figure 6

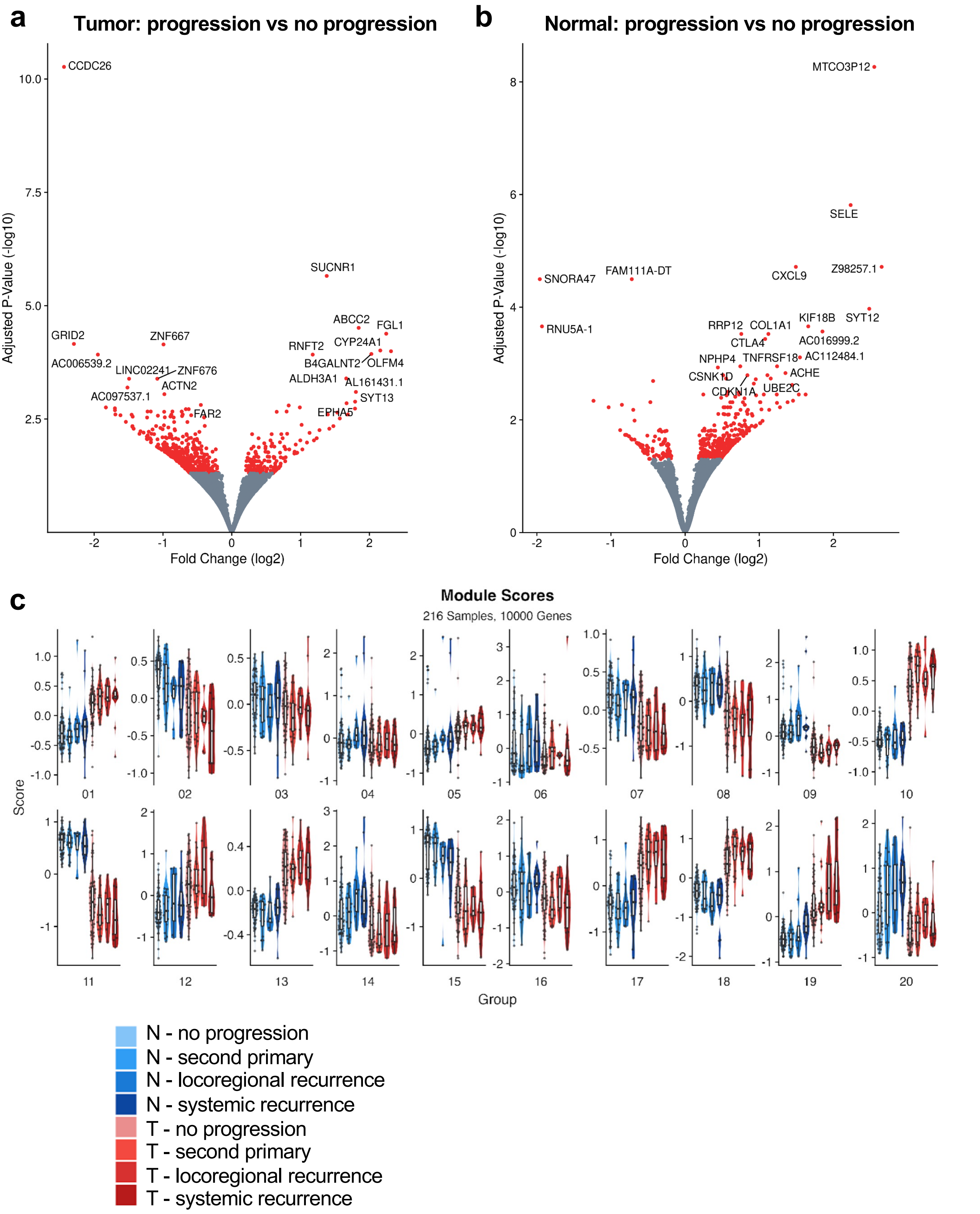

**Supplementary Figure 6: Transcriptomic signatures of disease progression in stage I lung adenocarcinoma.**

(a) Volcano plot representing the differential expression analysis comparing patients with progression vs no progression (tumor samples),

(b) Volcano plot representing the differential expression analysis comparing patients with progression vs no progression (normal lung samples),

(c) Boxplots comparing modules scores by progression type in tumor and normal tissue in each module.

### Supplementary Figure 7

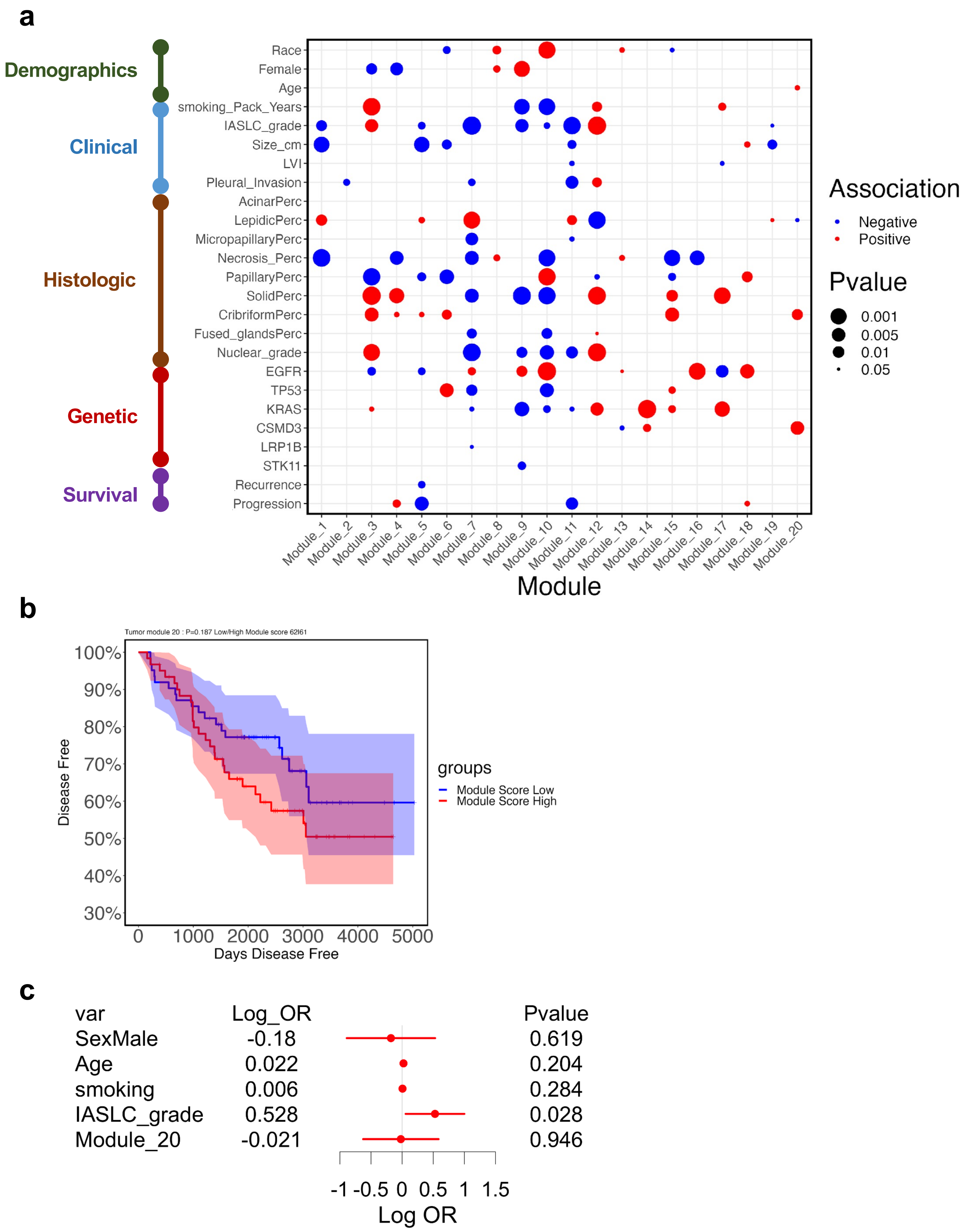

**Supplementary Figure 7: Association of module scores in tumor with different variables**

(a) Positive and negative associations of demographic, clinical, histologic, genetic and outcomes with module scores in tumor
(b) K-M PFS curve for patients with high and low module 20 scores in tumor
(c) Multivariate modeling of time-to-progression

### Supplementary Figure 8

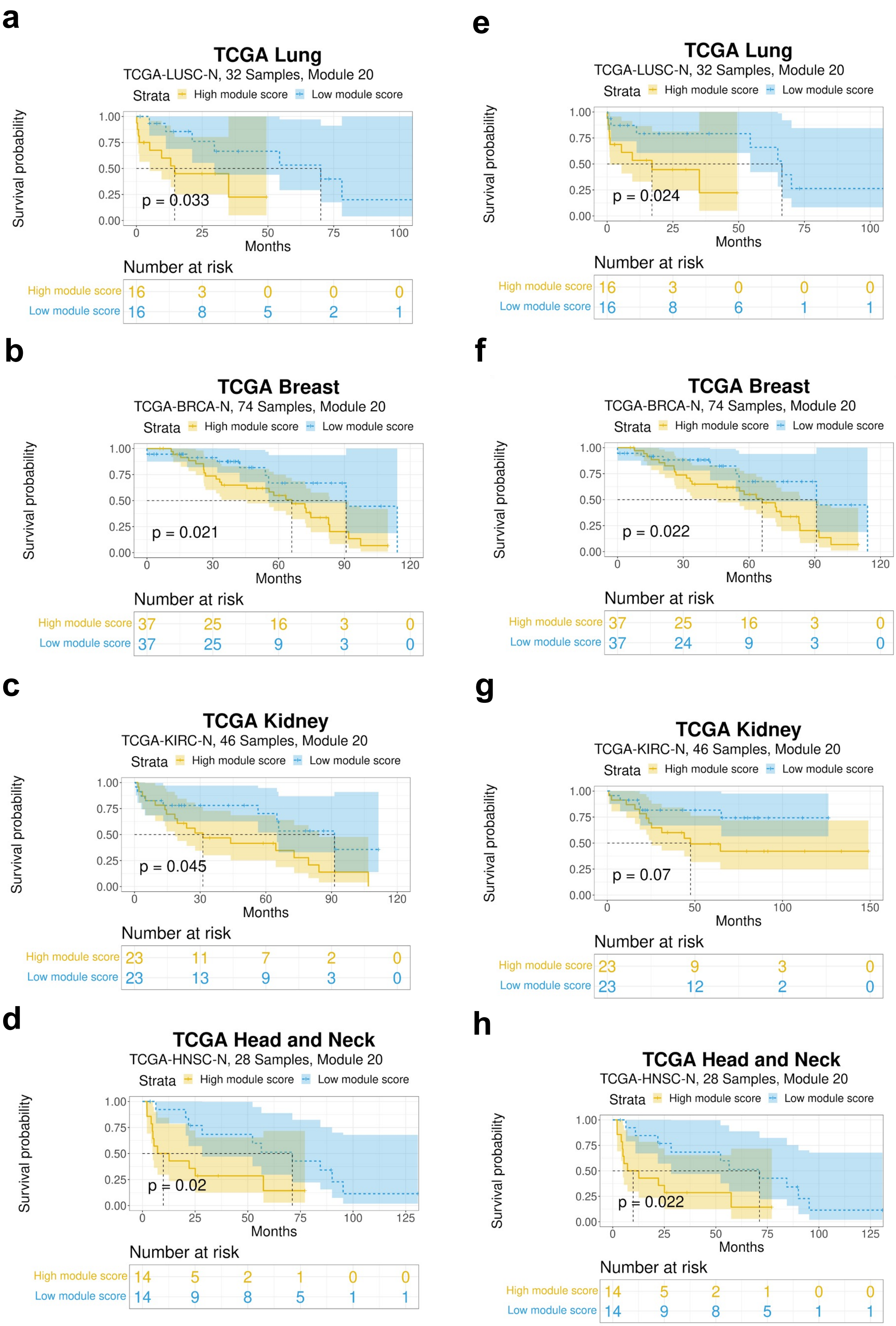

**Supplementary Figure 8: Progression-Free Survival (PFS) and Overall survival (OS) analysis of module 20 signature across TCGA cancer types**

(a) K-M PFS curves of patients with high and low module 20 scores in lung cancer
(b) K-M PFS curves of patients with high and low module 20 scores in breast cancer
(c) K-M PFS curves of patients with high and low module 20 scores in kidney cancer
(d) K-M PFS curves of patients with high and low module 20 scores in head and neck cancer
(e) K-M OS curves of patients with high and low module 20 scores in lung cancer
(f) K-M OS curves of patients with high and low module 20 scores in breast cancer
(g) K-M OS curves of patients with high and low module 20 scores in kidney cancer
(h) K-M OS curves of patients with high and low module 20 scores in head and neck cancer

### Supplementary Figure 9

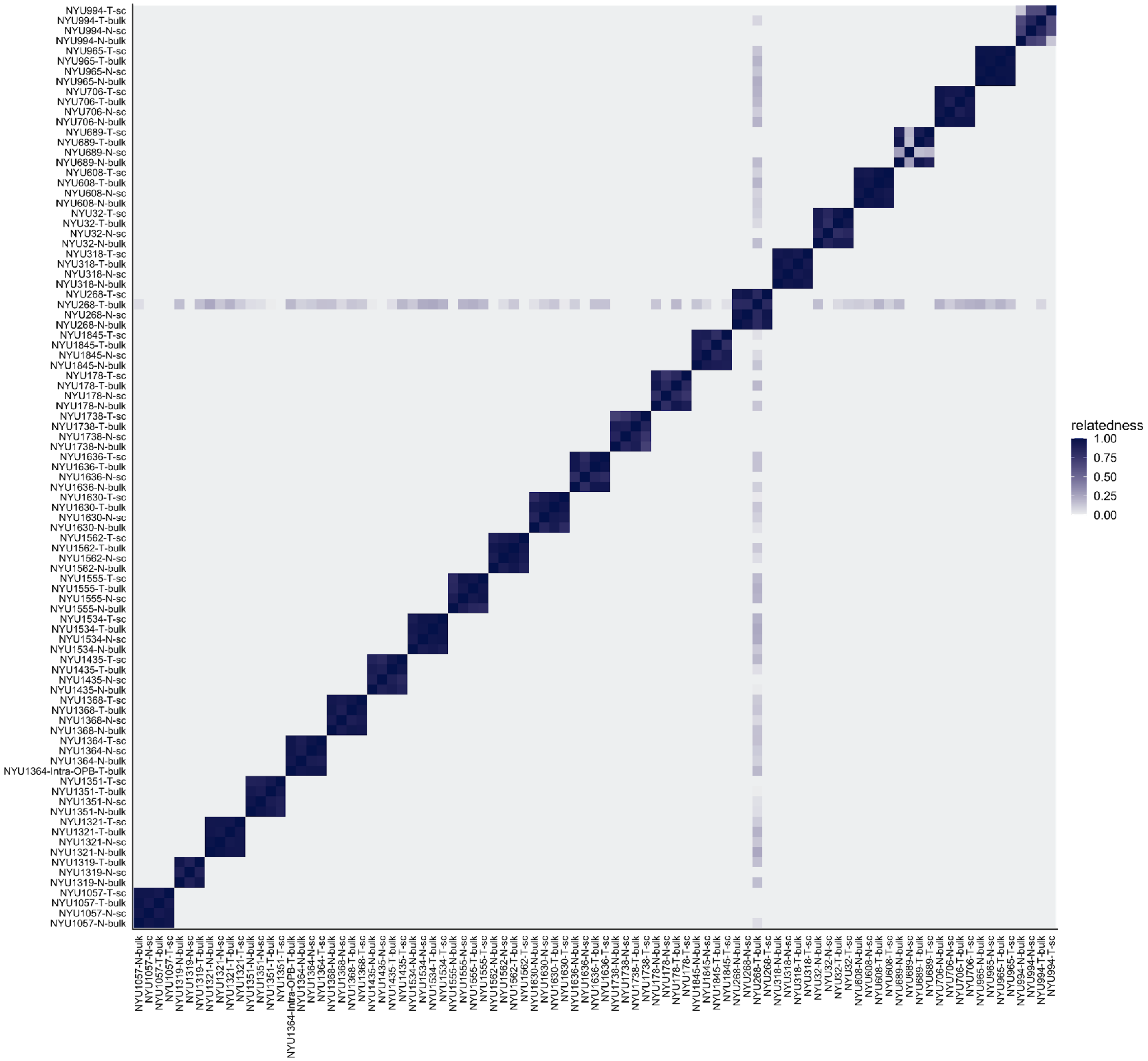

**Supplementary Figure 9: Genotype-based relatedness of single-nucleus RNA-seq tumor-normal samples to the corresponding bulk RNA-seq samples.**

### Supplementary Figure 10

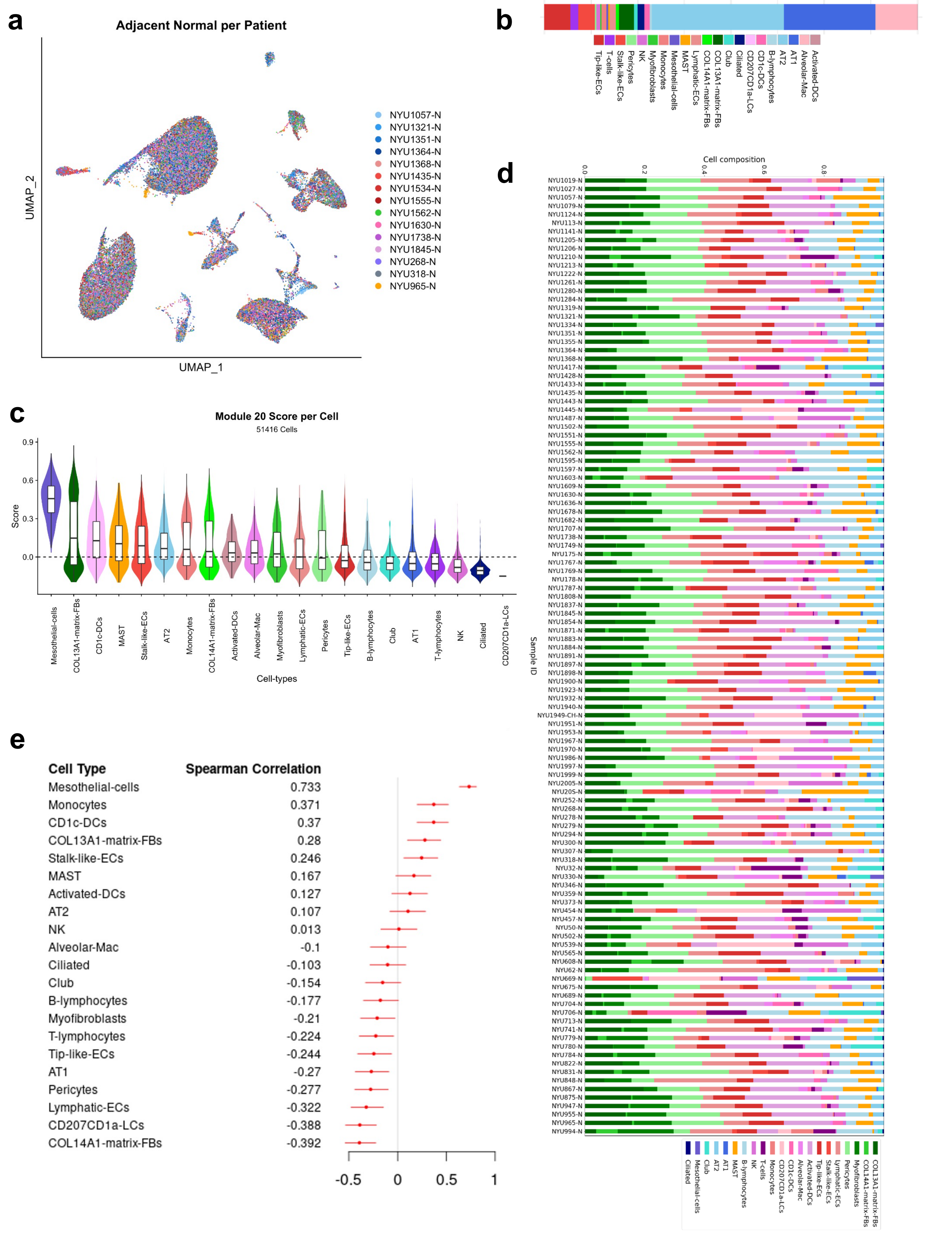

**Supplementary Figure 10: Single-nucleus RNA-seq analysis**

(a) UMAP visualization of all 51,428 adjacent normal cells, color-coded by patient
(b) Cell type abundance in the snRNA-seq dataset
(c) Per-cell module 20 score distributions grouped by cell type
(d) Cell type composition barcharts of BayesPrism-deconvoluted bulk RNA-seq tumor-adjacent normal samples
(e) Correlation of cell type composition with (bulk) module 20 scores across patients
